# PTEN mutant NSCLC require ATM to suppress pro-apoptotic signalling and evade radiotherapy

**DOI:** 10.1101/2021.07.24.453632

**Authors:** Thomas Fischer, Oliver Hartmann, Michaela Reissland, Cristian Prieto-Garcia, Kevin Klann, Christina Schülein-Völk, Bülent Polat, Elena Gerhard-Hartmann, Mathias Rosenfeldt, Christian Münch, Michael Flentje, Markus E. Diefenbacher

## Abstract

**Background:** Despite advances in treatment of patients with non-small cell lung cancer, carriers of certain genetic alterations are prone to failure. One such factor frequently mutated, is the tumor suppressor PTEN. These tumors are supposed to be more resistant to radiation, chemo- and immunotherapy.

**Methods:** Using CRISPR genome editing, we deleted PTEN in a human tracheal stem cell-like cell line as well generated primary murine NSCLC, proficient or deficient for *Pten*, *in vivo*. These models were used to verify the impact of PTEN loss *in vitro* and *in vivo* by immunohistochemical staining, western blot and RNA-Sequencing. Radiation sensitivity was assessed by colony formation and growth assays. To elucidate putative treatment options, identified via the molecular characterisation, PTEN pro- and deficient cells were treated with PI3K/mTOR/DNA-PK-inhibitor PI-103 or the ATM-inhibitors KU-60019 und AZD 1390. Changes in radiation sensitivity were assessed by colony-formation assay, FACS, western-blot, phospho-proteomic mass spectrometry and *ex vivo* lung slice cultures.

**Results:** We demonstrate that loss of PTEN led to altered expression of transcriptional programs which directly regulate therapy resistance, resulting in establishment of radiation resistance. While PTEN-deficient tumor cells were not dependent on DNA-PK for IR resistance nor activated ATR during IR, they showed a significant dependence for the DNA damage kinase ATM. Pharmacologic inhibition of ATM, via KU-60019 and AZD1390 at non-toxic doses, restored and even synergized with IR in PTEN-deficient human and murine NSCLC cells as well in a multicellular organotypic *ex vivo* tumor model.

**Conclusion:** PTEN tumors are addicted to ATM to detect and repair radiation induced DNA damage. This creates an exploitable bottleneck. At least *in cellulo* and *ex vivo* we show that low concentration of ATM inhibitor is able to synergise with IR to treat PTEN-deficient tumors in genetically well-defined IR resistant lung cancer models.

## MATERIAL AND METHODS

### Cell lines

Human BEAS-2B and HEK 293T cell lines was obtained from ATCC. Cells were maintained in high-glucose DMEM (Sigma Aldrich) supplemented with 10% FBS (Capricorn Scientific) and 1% Pen-Strep (Sigma Aldrich) 1% Glutamin (Sigma Aldrich) at 37°C in 5% CO_2_ on 10 cm dishes (Greiner Bioscience). For cell detachment Trypsin (Sigma Aldrich) was used. All the cells were maintained in culture for 15 passages as maximum to maintain cell identity. Cells were routinely tested for mycoplasma via PCR. The reagents were dissolved in Dimethyl sulfoxide (DMSO) in specified concentrations and added to the cells.

### DNA transfection and infection

For DNA transfection, a mix of 2,5 μg plasmid DNA, 200 μl free medium and 5 μl PEI was added into the 6-well dish medium (60% confluence), after 6 h incubation at 37°C the medium was changed to full supplemented medium. For DNA infection retroviruses or lentiviruses (MOI=10) were added to the cell medium in the presence of polybrene (5μg/ml) and incubated at 37°C for 72 h. After incubation, infected cells were selected with 2 μg/ml puromycin for 72 h or 20 μg/ml blasticidin for 10 days.

### X-ray irradiation

Irradiation was performed at room temperature using a 6 MV Siemens linear accelerator (Siemens, Concord, CA) at a dose rate of 9,5 Gy/min.

### Colony forming

Dependent on the experiment cells were treated with two different protocols. With the direct seeding protocol exponential growing cells were seeded to 10 cm dishes in adequate amount to be 50-80% confluent next day. Cells were trypsinized, counted and diluted. The dilution was dispensed into different vials and cells were irradiated in suspension. Cells were directly seeded in adequate amounts into 10 cm plates to obtain 100-400 colonies per dish. With the re-seeding protocol exponential growing cells were seeded to 10 cm dishes in adequate amount to be 25-30% confluent next day. The attached cells were treated with different substances or DMSO as a control. 3 h after treatment cells were irradiated with 0, 2, 3, 5, 7, 8 Gy and cultured for 24 h, then cells were trypsinized, counted and re-seeded in adequate amounts into 10 cm plates to obtain 100-400 colonies per dish. For both protocols KP and KPP cells formed colonies after 6 days, BEAS-2B cells formed colonies after 10-11 days. Cells were fixed with ice cold 25% acidic acid in methanol and stained with 0,5% crystal violet. Colonies were count manually. Only colonies containing at least 50 cells were scored. Surviving fractions were calculated by dividing the plating efficiency for the specified dose divided by the plating efficiency of untreated cells. Radiation treatment survival curves were fitted to the linear-quadratic model formula S= exp[−αD-βD^2^] (S=survival fraction; D=radiation dose; α and β fitted parameters). Curves were fitted and blotted using a non-linear regression and analysed with OriginPro (OriginPro, 2020, OriginLab Corporation, Northampton, MA, USA). Mean survival fractions at 2 Gy (SF2) and 4 Gy (SF4) were also obtained for each cell model and each substance and used to calculate the radiation enhancement ratio at 2 Gy (RER_2Gy_) and 4 Gy (RER_4Gy_) RER greater than 1 indicates enhancement of radiosensitivity, RER below the value of 1 indicates a radio resistance effect. Similarly, the radiation dose with 25% (D_25_) and 50% (D_50_) survival under different conditions was calculated to obtain the dose enhancement ratio (DER_25_ and DER_50_) that is calculated by dividing D_25_ without substance treatment by D_25_ with substance treatment, respectively D_50_ without substance treatment by D_50_ with substance. DER greater than 1 indicates a radio sensitising effect, a DER below the value of 1 indicates a radio protecting effect. Plating efficiency was calculated by dividing the number of colonies by the number of seeded cells. All calculated parameters are listed in supplementary table 1 (Table S1)

### Immunological methods

Cells were lysed in RIPA lysis buffer (20 mM Tris-HCl pH 7.5, 150 mM NaCl, 1 mM Na2EDTA, 1 mM EGTA, 1% NP-40 and 1% sodium deoxycholate), containing proteinase inhibitor an phosphatase inhibitor (1/100; Bimake) by sonication using Branson Sonifier 150 with a duty cycle at 25%, output control set on level 2 and the timer set to 15 s. Protein concentration was quantified using Bradford assay (Biorad). After mixing of Bradford reagent with 2 μl of sample, the photometer was used to normalize the protein amounts with a previously performed bovine serum albumin (BSA) standard curve. The quantified protein (40-80 μg) was heated in 4x sample buffer (Thermo Fisher) and 10% sample reducing agent (Thermo Fisher) for 10 min at 70°C and separated on 4-12% Bis/Tris-gels or 3-8% Tris/Acetat-Gels (Thermo Fisher). After separation, protein was transferred to nitrocellulose membrane (Thermo Fisher) in transfer buffer (Thermo Fisher) and then, incubated with blocking buffer (5% low fat milk powder in TBS and 0.1% Tween20) for 60 min at RT. After blocking, membranes were incubated with indicated Primary antibodies (1/1000 dilution in a buffer composed 5% low fat milk powder or 5% BSA in TBS and 0.1% Tween20) over night at 4°C. Secondary HRP coupled antibody (Dako 1/1000 dilution in a buffer composed 5% low fat milk powder or 5% BSA in TBS and 0.1% Tween20) were incubated for 2 h at 4°C. Membranes were incubated for 5 min in luminol-solution (250 mg luminol in 100 mM Tris pH 8,6) with 10% v/v cumarinic acid solution (1,1 g cumarinic acid in DMS0 and 0,1% v/v H_2_O_2_)at RT, then membranes were recorded with my ECL Imaging System. Analysis and quantifications of protein expression was performed using Image Studio software (Licor Sciences, Lincoln, NE, USA). Antibodies used for this publication are listed in supplemantary table 2 (Table S2)

### AnnexinV/DAPI staining

Cells growing as sub-confluent monolayers were pretreated with substance for 3 h before radiation with 0 Gy and 8 Gy. The cells were kept under standard conditions for normal cell growth. 24 h, 48 h, 72 h and 96 h after radiation cells were harvested with trysinization. Non irradiated cells, treated with camptothecin 5 μM (CPT) were harvested 48 h after treatment. Supernatant of cell culture dishes was pooled with trypsinized cells and pelleted by centrifugation. Further preparation for FACS measurement was following the protocol of the BioLegend APC Annexin V Apoptosis Detection Kit and DNA-staining with DAPI Reagent (25 μg/mL) (Biolegend, San Diego, CA, USA). 20 000 cells were assayed using a flow cytometer FACSCantoII (Becton Dickinson, San Jose, CA, USA). The output data presented as two-dimensional dot plot. Samples were analyzed using the Flowing software gating events to avoid debris, then dividing events in four quadrants. Flowing software was obtained from P. Terho (Turku Centre for Biotechnology, Turku, Finland). Column histograms and statistics were analyzed with Graphpad PRISM 8 (GraphPad Software, San Diego, California USA) and OriginPro. (OriginPro, 2020, OriginLab Corporation, Northampton, MA, USA).

### sgRNA design

sgRNAs were designed using the CRISPRtool (https://zlab.bio/guide-design-resources).

### AAV and lentivirus production and purification

Virus was packaged and synthetized in HEK 293T cells seeded in 15 cm-dishes. For AAV production, cells (70% confluence) were transfected with the plasmid of interest (10 μg), pHelper (15 μg) and pAAV-DJ or pAAV-2/8 (10 μg) using PEI (70 μg). After 96 h, the cells and medium of 3 dishes were transferred to a 50 ml Falcon tube together with 5 ml chloroform. Then, the mixture was shaken at 37°C for 60 min and NaCl (1 M) was added to the mixture. After NaCl is dissolved, the tubes were centrifuged at 20 000 × g at 4°C for 15 min and the chloroform layer was transferred to another Falcon tube together with 10% PEG8000. As soon as the PEG800 is dissolved, the mixture was incubated at 4°C overnight and pelleted at 20 000 × g at 4°C for 15 min. The pellet was resuspended in PBS with MgCl2 and 0.001% pluronic F68, then, the virus was purified using Chloroform and stored at − 80C. AAV viruses were titrated using Coomassie staining and RT-PCR using AAV-ITR sequence specific primers.

For Lentivirus production, HEK 293T cells (70% confluence) were transfected with the plasmid of interest (15 μg), pPAX (10 μg) and pPMD2 (10 μg) using PEI (70 μg). After 96h, the medium containing lentivirus was filtered and stored at −80°C.

### In vivo experiments and histology

All *in vivo* experiments were approved by the Regierung Unterfranken and the ethics committee under the license numbers 2532-2-362, 2532-2-367, 2532-2-374 and 2532-2-1003. The mouse strains used for this publication are listed. All animals are housed in standard cages in pathogen-free facilities on a 12 h light/dark cycle with *ad libitum* access to food and water. FELASA2014 guidelines were followed for animal maintenance.

Adult mice were anesthetized with Isoflurane and intratracheally intubated with 50 μl AAV virus (3 × 10^7^ PFU) as previously described (Prieto-Garcia et al. 2019). Viruses were quantified using Coomassie staining protocol^1^. Animals were sacrificed by cervical dislocation and lungs were fixed using 10% NBF. H&E, slides were de-paraffinized and rehydrated following the protocol: 2x 5 min. Xylene, 2x 3 min. EtOH (100%), 2x 3 min. EtOH (95%), 2x 3 min. EtOH (70%), 3 min. EtOH (50%) and 3 min. H_2_O. For all staining variants, slides were mounted with 200 μl of Mowiol® 40-88 covered up by a glass coverslip. IHC slides were recorded using Panoramic DESK scanner or using FSX100 microscopy system (Olympus) and analysed using Case Viewer software (3DHISTECH) and ImageJ.

### Primary murine lung cancer cell lines

In brief, at endpoint of experiment, tumor bearing mice were sacrificed and lung lobes excised. The tissue was briefly rinsed in PBS and transferred to PBS containing Petri dishes. By using a binocular, macroscopically detectable tumor lesions on the lung lobes were excised with a scissor and transferred to a test tube containing Collagenase I (100 U/ml in PBS). The tumor containing tissue was digested for 30 min at 37°C, and the reaction was stopped by addition of 10% FCS. The tissue/collagenase/FCS mixture was briefly spun in a benchtop centrifuge and the supernatant discarded. Digested tissue was re-suspended in 10% FCS (Capricorn) DMEM (Sigma Aldrich), Pen/Strep (Sigma Aldrich) and washed 3 times in 1 ml solution prior to plating in a 6 well tissue culture plate. During subsequent re-plating fibroblasts were counter-selected, by selective trypsinisation, and cell clusters with a homogenous morphology were clonally expanded. These clones were then subjected to further biochemical analysis and characterisation, including genotyping PCR, RNA-sequencing.

### Tumor area

FFPE fixed tissue sections from animals were de-parafinized and stained with haematoxylin and eosin (H&E). Each slide was scanned using a Roche Ventana DP200 slide scanner. To assess tumor area per animal, total lung area was measured by using the QuPath image analsyis tool. Subsequently, all tumor nodules were measured and the tumor surface calculated. Graph was generated using GraphPad Prism 8.

### Survival curves mouse

Upon intratracheal administration of AAV, animals were monitored on a daily basis. Whenever experimentally defined termination points were reached, such as 20% weight loss, animals were sacrificed by cervical dislocation and tissue samples collected. Graphs were generated using Prism Graphpad 8.

### RNA-sequencing

RNA sequencing was performed with Illumina NextSeq 500 as described previously^2^. RNA was isolated using ReliaPrep™ RNA Cell Miniprep System Promega kit, following the manufacturer’s instruction manual. mRNA was purified with NEBNext® Poly(A) mRNA Magnetic Isolation Module (NEB) and the library was generated using the NEBNext® UltraTM RNA Library Prep Kit for Illumina, following the manufacturer’s instructions).

### Sample preparation for mass spectrometry

Lysates of cells, solved from cell culture plates with cell scrapers pelleted and frozen at −80°C, were precipitated by methanol/chloroform and proteins resuspended in 8 M Urea/10 mM EPPS pH 8.2. Concentration of proteins was determined by Bradford assay and 300 μg of protein per samples was used for digestion. For digestion, the samples were diluted to 1 M Urea with 10 mM EPPS pH 8.2 and incubated overnight with 1:50 LysC (Wako Chemicals) and 1:100 Sequencing grade trypsin (Promega). Digests were acidified using TFA and tryptic peptides were purified by Oasis Prime HLB columns (30 mg, Waters). 80 μg peptides per sample were TMTpro labeled, and the mixing was normalized after a single injection measurement by LC-MS/MS to equimolar ratios for each channel. 100 μg of pooled peptides were dried for offline High pH Reverse phase fractionation by HPLC (whole cell proteome) and remaining ~1.1 mg of multiplexed peptides were used for phospho-peptide enrichment by High-Select Fe-NTA Phosphopeptide enrichment kit (Thermo Fisher) after manufacturer’s instructions. After enrichment, peptides were dried and resuspended in 70% acetonitrile/0.1% TFA and filtered through a C8 stage tip to remove contaminating Fe-NTA particles. Dried phospho-peptides then were fractionated on C18 (Empore) stage-tip. For fractionation C18 stagetips were washed with 100% acetonitrile twice, followed by equilibration with 0.1% TFA solution. Peptides were loaded in 0.1% TFA solution and washed with water. Elution was performed stepwise with different acetonitrile concentrations in 0.1% Triethylamine solution (5%, 7.5%, 10%, 12.5%, 15%, 17.5%, 20%, 22.5%, 25%, 27.5%, 30%, 50%). The resulting 12 fractions were concatenated into six fractions and dried for LC-MS.

Peptides were fractionated using a Dionex Ultimate 3000 analytical HPLC. 250 μg of pooled and purified TMT-labeled samples were resuspended in 10 mM ammonium-bicarbonate (ABC), 5% ACN, and separated on a 250 mm long C18 column (X-Bridge, 4.6 mm ID, 3.5 μm particle size; Waters) using a multistep gradient from 100% Solvent A (5% ACN, 10 mM ABC in water) to 60% Solvent B (90% ACN, 10 mM ABC in water) over 70 min. Eluting peptides were collected every 45 s into a total of 96 fractions, which were cross-concatenated into 24 fractions and dried for further processing.

#### Liquid chromatography mass spectrometry

All mass spectrometry data was acquired in centroid mode on an Orbitrap Fusion Lumos mass spectrometer hyphenated to an easy-nLC 1200 nano HPLC system using a nanoFlex ion source (ThermoFisher Scientific) applying a spray voltage of 2.6 kV with the transfer tube heated to 300°C and a funnel RF of 30%. Internal mass calibration was enabled (lock mass 445.12003 m/z). Peptides were separated on a self-made, 32 cm long, 75 μm ID fused-silica column, packed in house with 1.9 μm C18 particles (ReproSil-Pur, Dr. Maisch) and heated to 50°C using an integrated column oven (Sonation). HPLC solvents consisted of 0.1% Formic acid in water (Buffer A) and 0.1% Formic acid, 80% acetonitrile in water (Buffer B).

For total proteome analysis, a synchronous precursor selection (SPS) multi-notch MS3 method was used in order to minimize ratio compression as previously described. Individual peptide fractions were eluted by a non-linear gradient from 7 to 40% B over 90 min followed by a step-wise increase to 95% B in 6 min which was held for another 9 min. Full scan MS spectra (350-1400 m/z) were acquired with a resolution of 120,000 at m/z 200, maximum injection time of 100 ms and AGC target value of 4 × 105. The most intense precursors with a charge state between 2 and 6 per full scan were selected for fragmentation and isolated with a quadrupole isolation window of 0.7 Th and a cycle time of 1.5 s. MS2 scans were performed in the Ion trap (Turbo) using a maximum injection time of 50 ms, AGC target value of 1.5 × 104 and fragmented using CID with a normalized collision energy (NCE) of 35%. SPS-MS3 scans for quantification were performed on the 10 most intense MS2 fragment ions with an isolation window of 0.7 Th (MS) and 2 m/z (MS2). Ions were fragmented using HCD with an NCE of 65% and analyzed in the Orbitrap with a resolution of 50,000 at m/z 200, scan range of 110-500 m/z, AGC target value of 1.5 × 105 and a maximum injection time of 120 ms. Repeated sequencing of already acquired precursors was limited by setting a dynamic exclusion of 45 seconds and 7 ppm and advanced peak determination was deactivated.

For phosphopeptide analysis, each peptide fraction was eluted by a linear gradient from 5 to 32% B over 120 min followed by a step-wise increase to 95% B in 8 min which was held for another 7 min. Full scan MS spectra (350-1400 m/z) were acquired with a resolution of 120,000 at m/z 200, maximum injection time of 100 ms and AGC target value of 4 × 105. The most intense precursors per full scan with a charge state between 2 and 5 were selected for fragmentation, isolated with a quadrupole isolation window of 0.7 Th and fragmented via HCD applying an NCE of 38% with an overall cycle time of 1.5 s. MS2 scans were performed in the Orbitrap using a resolution of 50,000 at m/z 200, maximum injection time of 86ms and AGC target value of 1 × 105. Repeated sequencing of already acquired precursors was limited by setting a dynamic exclusion of 60 s and 7 ppm and advanced peak determination was deactivated.

### QUANTIFICATION AND STATISTICAL ANALYSIS

#### RNA-sequencing analysis

Fastq files were generated using Illuminas base calling software GenerateFASTQ v1.1.0.64 and overall sequencing quality was analyzed using the FastQC script. Reads were aligned to the human genome (hg19) using Tophat v2.1.1^3^ and Bowtie2 v2.3.2^4^ and samples were normalised to the number of mapped reads in the smallest sample. For differential gene expression analysis, reads per gene (Ensembl gene database) were counted with the “summarizeOverlaps” function from the R package “GenomicAlignments” using the “union”-mode and non- or weakly expressed genes were removed (mean read count over all samples <1). Differentially expressed genes were called using edgeR^5^ and resulting p-values were corrected for multiple testing by false discovery rate (FDR) calculations. GSEA analyses were done with signal2Noise metric and 1000 permutations. Reactome analysis were performed with PANTHER^6^ using the “Statistical overrepresentation test” tool with default settings. Genes were considered significantly downregulated for Reactome analysis when: Log2FC>0.75 and FDR p-value<0.05.

#### Analysis of publicly available data

All publicly available data and software used for this publication are listed (please see Star Methods). Oncoprints were generated using cBioportal^7, 8^. Briefly, Oncoprints generates graphical representations of genomic alterations, somatic mutations, copy number alterations and mRNA expression changes. TCGA data was used for the different analysis. Data were obtained using UCSC Xena. Data was downloaded as log2 (norm_count+1)

Kaplan-Meier curves were estimated with the KM-plotter^9^, cBioportal^7^ and R2: Genomics Analysis and Visualization Platform (http://r2.amc.nl). The KM-plotter was used to analyse overall survival of lung cancer patients (Figure 1 and S1) based on gene expression data from microarrays obtained from GEO, caBIG and TCGA

**Figure 1:**
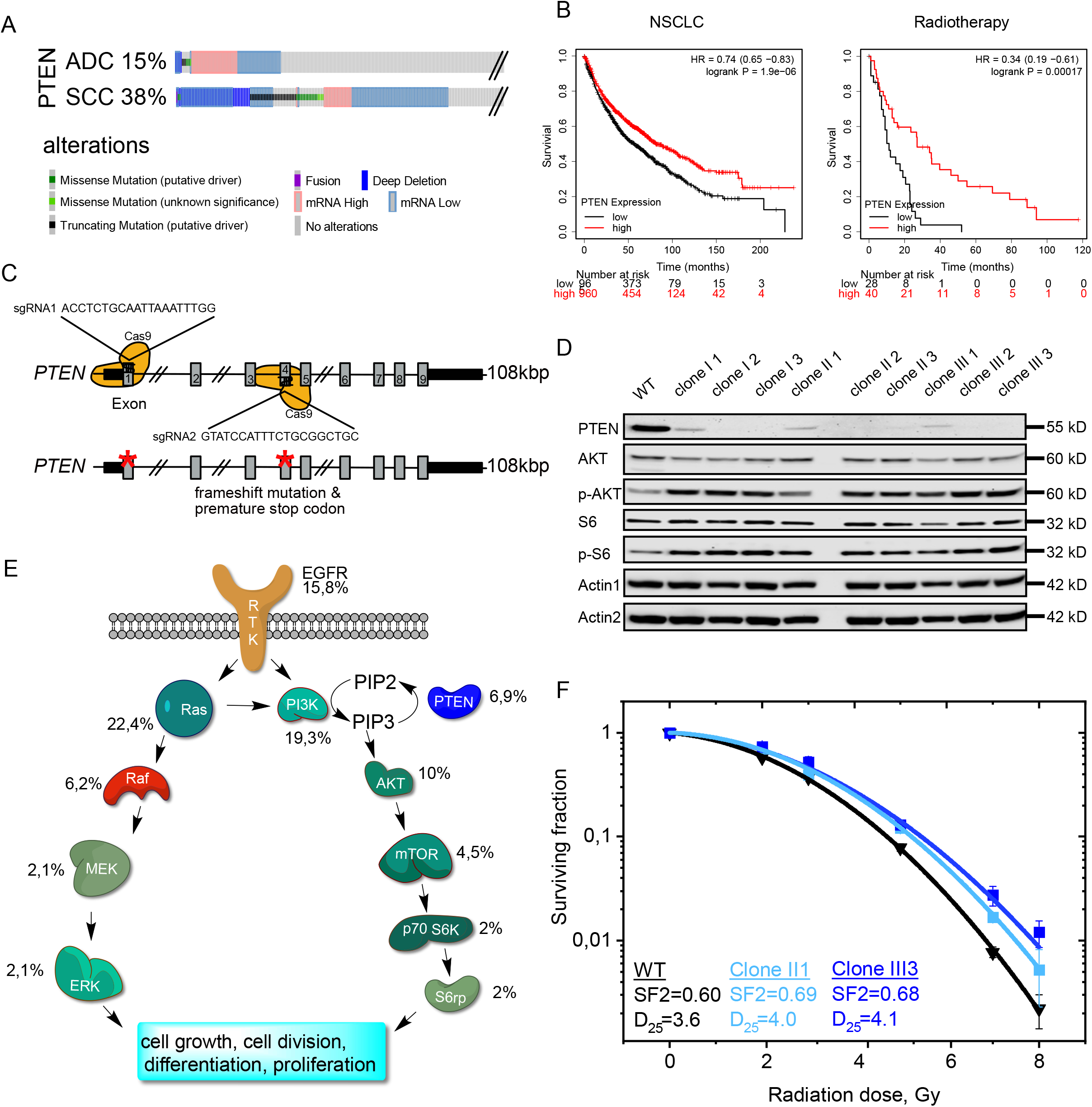
PTEN alterations; impact on pathways and radiation resistance A) PTEN alteration frequency of NSCLC from cBioportal, https://www.cbioportal.org/ Analysis of Lung Squamous Cell Carcinoma (SCC) and Adenocarcinoma (ADC). B) Kaplan-Meier Plot of lung cancer patient survival stratified by PTEN (204054_at) expression. Median survival in the ADC cohort of the low PTEN expression 61.3 months, of high expression 175 months. Median survival in the SCC cohort of the low PTEN expression 42 months, of high expression 72.3 months. The p-value was calculated using a logrank test. HR: hazard ratio. Generated with the online tool https://kmplot.com. C) Schematic representation of the CRISPR/Cas9 genome editing strategy to delete PTEN in the human lung cell line BEAS-2B targeting exon 1 and exon 4. D) Immunoblot of virus transfected, blasticidin selected and clonogenic isolated BEAS-2B cells, generated with the described method (Supp. Figure 2B). Control: WT: Epithelial transformed BEAS-2B PTEN^wt^ cells. Actin as loading control. E) Receptor-tyrosine-kinase signaling cascade of the MAPK-pathway and PI3K/Akt pathway. Numbers next to the Enzymes show the percentage of genetic alteration of the coding genes. Data generated with the free online tool https://www.cbioportal.org. F) Colony formation assay BEAS-2B clone II1 (PTEN^*hetero*^, light blue) and III3 (PTEN^*homo*^, blue) compared to vector control (black). SF 2: Surviving fraction at 2 Gy. D_25_: Dose in Gy with 25% survival. Error bars: Standard deviation. n=3. Also see Supplementary Figure S1.

#### Mass spectrometry data analysis

Raw files were analyzed using Proteome Discoverer (PD) 2.4 software (ThermoFisher Scientific). Spectra were selected using default settings and database searches performed using SequestHT node in PD. Database searches were performed against trypsin digested Mus musculus SwissProt database containing one sequence per gene without isoforms. Static modifications were set as TMTpro at the N-terminus and lysines and carbamidomethyl at cysteine residues. Search was performed using Sequest HT taking the following dynamic modifications into account: Oxidation (M), Phospho (S,T,Y), Met-loss (Protein N-terminus), Acetyl (Protein N-terminus) and Met-loss acetyl (Protein N-terminus). For whole cell proteomics, the same settings were used except phosphorylation was not allowed as dynamic modification. For phospho-proteomics all peptide groups were normalized by summed intensity normalization and then analyzed on peptide level^10^. For whole cell proteomics normalized PSMs were summed for each accession and data exported for further use. For proteomics analysis, significance was assessed via a two-sided unpaired students t-test with equal variance assumed. For pathway analysis, Protein/Peptide lists were filtered as indicated and a STRING network created in Cytoscape. For the resulting network a pathway enrichment analysis was performed using the STRING App Cytoscape plugin. For network views of enrichments, the Reactome pathways were filtered for a FDR < 0.001 and loaded into the Enrichment Map 3 plugin for Cytoscape to create visualization. Gene sets for visualization purposes were downloaded from the molecular signature gene set database (https://www.gsea-msigdb.org/) on 02-21-2021. Result files were filtered for the included genes to create pathway specific visualizations.

### DATA AND SOFTWARE AVAILABILITY

Raw data is available via Mendeley Data: doi: RNA-sequencing data is available at the Gene Expression Omnibus under the accession number GEO:

### Contact for reagent and resource sharing

Further information and requests for resources and reagents should be directed to and will be fulfilled by the Lead Contact, Markus E. Diefenbacher.

## Introduction

Lung cancer is the most common cancer worldwide, claiming 1.76 million lives in 2018 alone (WHO cancer statistics 2018). This is exceeding total numbers of colon, breast and prostate cancer combined^11–14^. In the past decade, with the advent of targeted and immune-checkpoint blockade therapy, major improvements in treatment response of advanced NSCLC (non-small cell lung cancer) were achieved^15^. Targeted therapies are predominantly validated in the treatment of late stage patients (UICC stage IV)^16^. Other patients (UICC stage I, II and III)^17^, rarely benefit from these combinatorial treatments and survival rates have only marginally improved, with many patients still succumb to lung cancer within five years^18^. Furthermore, not all patients benefit equally from these novel therapeutic approaches^19–21^. Genetic analysis of tumor samples by Next Generation Sequencing (NGS) from treatment resistant patients highlighted that several genetic alterations can contribute to therapy resistance and reduced patient survival e.g. *KRAS, STK11, KEAP1* and the phosphatase and tensin homologue (*PTEN*)^22–24^.

PTEN was initially described as a phosphatase involved in the homeostatic maintenance of the phosphatidylinositol-3-kinase/protein kinase B (PI3K/AKT) cascade leading to suppression of phospho-AKT^25^. It functions as a tumor suppressor via affecting cell cycle progression, inhibition of cell death, transcription, translation, stimulation of angiogenesis, and maintenance of stem cell self-properties via mTOR signalling^26^. NGS of tumor samples revealed that PTEN is frequently deleted or mutated in a variety of tumors^12^, including NSCLC (Adenocarcinoma and Squamous cell carcinoma)^13, 14^. PTEN itself, as a tumor suppressor, is not a direct target for cancer therapy, but can serve as a prognostic marker^27^. Mutations in PTEN result in resistance towards *‘standard of care’* therapies, such as radiotherapy and chemotherapy, by hyperactivation of the AKT pathway^28^. Additionally, loss of PTEN limits the employment of personalized therapy, as it is blunting therapeutic responses relying on immune checkpoint blockade and drives resistance to established targeted therapies like EGFR antagonists^29, 30^. Nuclear PTEN is involved in the control of essential biological processes, such as maintenance of genome integrity^31^, APC/C-CDH1-dependent PLK and AURK degradation^32^, chromatin remodelling^33^ and double strand break repair^34^.

DNA damage inducing therapies, such as ionizing radiation (IR), rely on the inability of tumor cells to efficiently clear all damage, while wild type cells undergo cell cycle arrest to gain sufficient time to repair^35, 36^. Here, DNA damage sensing enzymes, such as DNA-PK, ATR and ATM are key players and dictate the route taken for repair of the damaged DNA^37, 38^. Ataxia telangiectasia mutated kinase (ATM) is the prime sensor of double strand breaks induced by ionizing radiation^38^. It is required for downstream activation of SMC1, CHEK2, RAD50-MRE11 and BRCA1 signalling cascades, thereby contributing to radiation resistance and cell cycle checkpoint progression and arrest^39^. An alternative source of ATM activation is the induction of reactive oxygen species, a by-product of IR therapy^40^. Previous reports also highlighted a deregulation of ATM in PTEN mutant tumors, suggesting that the ATM-PTEN axis is of therapeutic value for certain cancers^41, 42^. Together, these data argue that inhibition of DNA damage sensors may restore therapy responses in PTEN mutant NSCLC and suggest that this strategy may have therapeutic efficacy in lung cancer.

## Results

### Alterations in PTEN affect patient survival and radiation therapy outcome in NSCLC

To assess the mutational as well the expression status of PTEN in human malignancy, we analysed public available patient data. Alterations in PTEN were frequently observed in lung cancer, both in adenocarcinoma (ADC) and squamous cell carcinoma (SCC), ranging between 15% and 38%, respectively (Figure 1A). PTEN is frequently altered in invasive tumors and reduced expression or mutation correlate with overall shortened patient survival (Figure 1B and S1A). Irrespective of NSCLC subtype, patient data suggest that PTEN loss is a direct prognostic marker for shorter survival, including tumor mutational burden (TMB) low patients, which are otherwise not amenable to immunotherapy and treated with chemotherapy (Figure S1A). Not only do *PTEN^mutant/low^* patients have an overall shortened life expectancy; when treated with radiotherapy alone, they have a poorer overall survival (p=0,00017) compared to a *PTEN^high^* patient cohort (Figure 1B). These data demonstrate that reduced expression or mutation of *PTEN* is a frequent event and significantly correlates with poor patient survival and treatment failure.

### Radiation sensitivity is PTEN-dosage dependent

Next, we investigated if loss of *PTEN* contributes to radiotherapy resistance. Instead of using classic human lung cancer cell lines with a high mutational burden, we utilized the human lung tracheal stem cell like cell line BEAS-2B. By using differentiated BEAS-2B cells we were on the one hand able to avoid putative mutations contributing to IR resistance, on the other hand we could mimic tumors with low TMB and worse patient survival outcome, with bigger need for successful treatment options. Deletion of PTEN in BEAS-2B was achieved by simultaneous CRISPR/Cas9 mediated gene editing of *PTEN* exon 1 and exon 4 (Figure 1C). BEAS-2B cells were lentivirally infected and upon Blasticidin selection, individual clones were analysed (Figure S1B). CRISPR/Cas9 mediated genome editing yielded heterozygous as well as homozygous deletions of *PTEN*, as seen by immunoblotting against endogenous PTEN (Figure 1D). As previously reported, loss of PTEN led to enhanced phosphorylation of AKT and its downstream target, S6 kinase, as seen by western blotting (Figure 1D and E). It is noteworthy that heterozygous loss of PTEN was sufficient to activate downstream phosphorylation cascades. Generated clones were expanded and subjected to a single dose of ionizing radiation of 2, 3, 5, 7 or 8 Gy, respectively. Upon irradiation, cells were directly re-seeded from suspension and colony formation capacity was assessed by crystal violet staining (Figure 1F, S1C and D). While *PTEN^wt^* BEAS-2B demonstrated an IR dosage dependent ability to form colonies, clones depleted for PTEN, *PTEN^clone-II(het)^* and *PTEN^clone-III(homo)^*, tolerated higher doses of IR, indicating that PTEN loss contributes to IR resistance (Figure 1F and S1D). Since mutations in PTEN co-occur with mutations in oncogenic drivers, we tested the impact of aberrant MAPK signalling on IR resistance (Figure 1E and S1F). By retroviral transduction, a mutant form of BRAF, BRAF^V600E^, was introduced in the clonal lines *PTEN^wt^* and *PTEN^clone-III3(homo)^* BEAS-2B. Overexpression of the mutant *V600E* variant of *BRAF* was detectable and resulted in the downstream activation of the MAPK pathway, as seen by phosphorylation of MEK (Figure S1E). Oncogenic BRAF^V600E^ did not alter the radiation sensitivity of *PTEN^wt^* BEAS-2B nor affected the relative resistance of *PTEN^homo^* BEAS-2B (Figure S1F).

These data demonstrate that genetic loss or mutation of PTEN is sufficient to establish IR resistance in the human non-oncogenic cell line BEAS-2B.

### Loss of Pten cooperates with mutant Tp53 and KRas^G12D^ in murine NSCLC in vivo and diminishes radiationsensitivity ex vivo

To investigate if the observed IR resistance is limited to “stable” cell lines or is a ‘hardwired’ feature of PTEN mutant tumors, we used CRISPR-mediated NSCLC mouse models driven by either mutations of Tp53 and KRas (KP: *KRas^G12D^,Tp53^mut^*) and studied the impact of an additional deletion of Pten (KPP : *KRas^G12D^:Tp53^mut^:Pten^mut^*). Constitutive Cas9 expressing mice were infected via intratracheal administration with an adeno-associated virus (AAV), packaged with the ubiquitous rep/cap 2/DJ^43^. 12 weeks post infection tumor burden and viability were assessed (Figure S2A). While *KP* mice developed tumors resulting in an overall transformation of around 16% of lung tissue, additional loss of *Pten* (KPP) proven by immunohistochemistry greatly enhanced the tumor area to 80% (Figure 2A, B and C). Additionally, as reported for patients, loss of *Pten* negatively affected survival (Figure 2D). While *KP* mice survived 12 weeks without showing physiological effects caused by their tumor burden, *KPP* mice required premature termination due to onset of various symptoms, such as weight loss/cachexia and increased breathing frequency (Figure 2D). Next, we established tumor cell lines from various animals by ectopic dissection of tumors and subsequent culture in standard medium (DMEM/10%serum/5%Pen/Strep)^44^. The genetic status of four established cell lines (KP5 and KP6; *KRas^G12D^:Tp53^mut^*; KPP4 and KPP8; *KRas^G12D^:Tp53^mut^:Pten^mut^*) was confirmed by polymerase chain reaction of genomic DNA derived from tumor cells. Loss of *Pten* and activation of the downstream pathway was further confirmed by immunoblotting and immunohistochemistry, showing increased phosphorylation of AKT in *KPP* when compared to *KP* tumors and primary tumor cell lines (Figure 2B, E and S2B and C). Exposure to IR significantly reduced the capacity of *KP* cells to establish colonies (Figure 2F). *KPP* tolerated higher doses of ionizing radiation compared to Pten wild type cells, reproducing the results obtained in the human cell line BEAS-2B *PTEN^homo^* (Figure 1F and 2F). Immunoblot analysis of pathway components of the PI3K and MAPK pathway of KP and KPP clones demonstrated that cells depleted of *Pten* maintained elevated expression of EGFR and phosphorylated AKT during ionizing irradiation, while other components of the pathway were not affected (Figure S2B and C).

**Figure 2:**
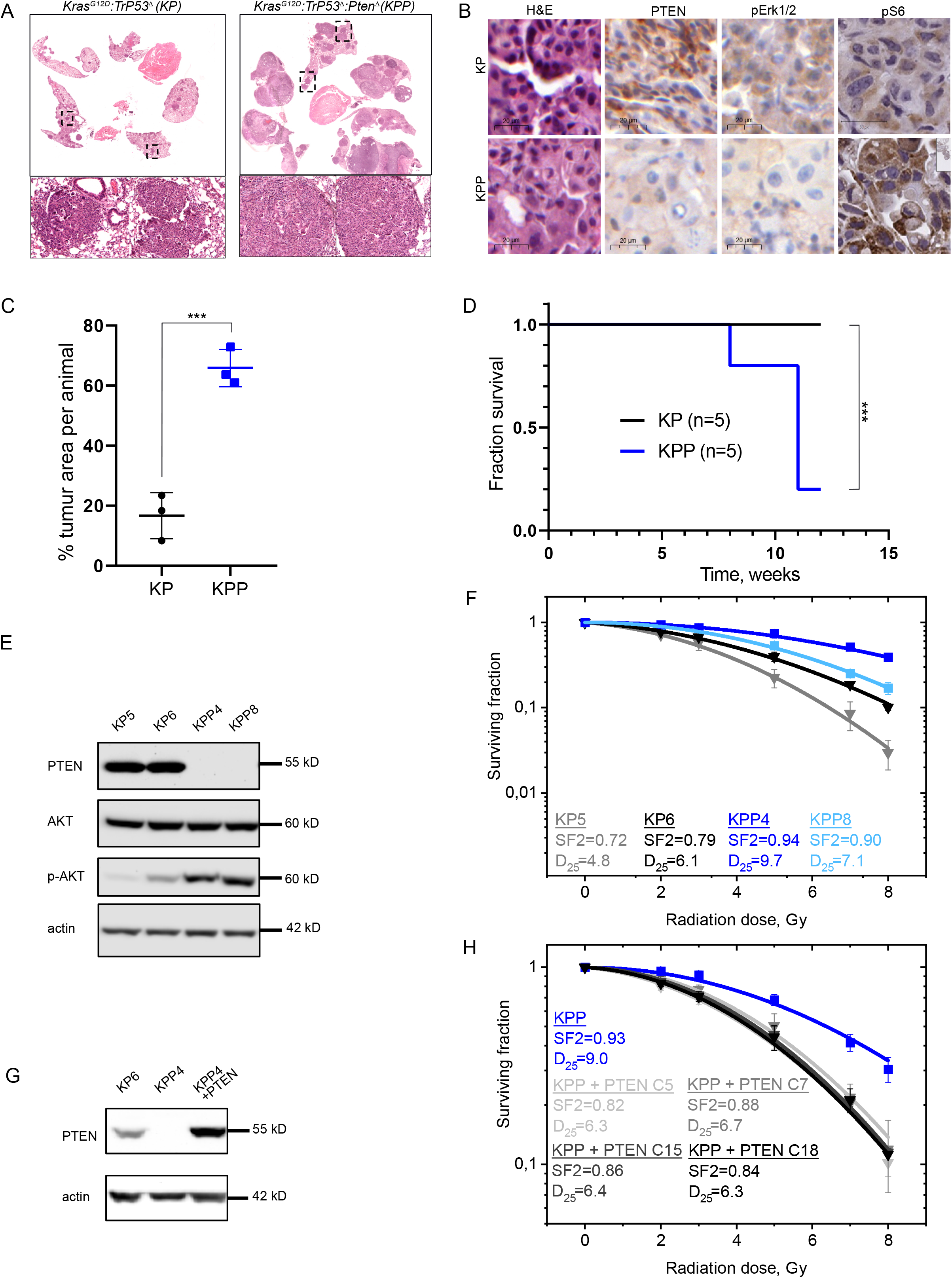
Generating and characterizing murine PTEN deficient tumor cell lines A) Representative haematoxylin and eosin (H&E) staining of tumor bearing animals 12 weeks post intratracheal infection. On the left KP (*KRas^G12D^:Tp53^mut^*) on the right KPP (*KRas^G12D^ :Tp53^mut^:Pten^mut^*). Boxes indicate upper highlighted tumor areas. B) Representative haematoxylin and eosin (H&E) and immunohistochemical DAB staining (PTEN, p-ERK1/2 and p-S6) of tumor bearing animals 12 weeks post intratracheal infection. on the upper part KP (*KRas^G12D^:Tp53^mut^*) on the lower part KPP (*KRas^G12D^ :Tp53^mut^:Pten^mut^*). C) Quantification of % tumor area (normalized to total lung area) in KP (black) and KPP (blue) animals. n=3. D) Kaplan-Meier survival curves comparing KP (black; n=5) and KPP (blue, n=5) animals upon AAV intratracheal infection. E) Immunoblot of endogenous (phospho-)AKT of two representative generated cell lines from different mice. KP5 and KP6 (*KRas^G12D^:Tp53^mu^*), KPP4 and KPP8 (*KRas^G12D^:Tp53^mut^:Pten^mut^*). Actin as loading control. n=3. F) Colony formation assay KP5 (gray), KP6 (black), KPP4 (blue) and KPP8 (light blue). SF 2: Surviving fraction at 2 Gy. D_25_: Dose in Gy with 25% survival. Error bars: Standard deviation. n=3. G) Immunoblot against PTEN/Pten of KP6 and lentivirally transduced, either GFP or human PTEN cDNA overexpressing KPP4 cells after Puromycin selection. Actin as loading control. n=3. H) Colony formation assay KPP4 (blue) and PTEN reconstituted KPP4 clones (C5, C7, C15 and C18; gray to black; Supp. Figure 3D) after clonogenic isolation. SF 2: Surviving fraction at 2 Gy. D_25_: Dose in Gy with 25% survival. Error bars: Standard deviation. n=3. Also see Supplementary Figure S2.

For proof of principle, we reconstituted the radiation resistant clone KPP4 with a human full length wild type PTEN cDNA, using lentiviral transduction (Figure 2G and S2D). PTEN expression was confirmed by immunoblotting against Pten/PTEN. Radiation dose response of several reconstituted clones was measured using colony survival. All clones expressing human PTEN showed enhanced sensitivity towards IR when compared to the parental PTEN^mut^ clone (Figure 2H).

Our data demonstrate that loss of *Pten* synergises with loss of *Trp53* and oncogenic *KRas* in NSCLC and accelerated tumor growth in the mouse lung. Cell lines derived from these tumors and lacking *Pten* showed decreased radiation sensitivity.

### Loss of Pten alters DNA damage signalling pathways

To understand how Pten loss affects overall gene expression, and if these changes could account for the IR resistance of *PTEN^mutant^* cells, we performed transcriptomic analysis by RNA sequencing of KP6 and KPP4 (from here on KP and KPP). While KP and KPP derived tumor cells share a high degree of commonly regulated genes (Spearman R=0.9122, Figure 3A), KPP cells upregulated 2441 distinct genes when compared to KP (Figure 3B). Gene set enrichment analysis (GSEA) showed that KP cells are predominantly driven by the KRas pathway, while cells mutant for *PTEN* altered the transcriptional profile towards the AKT1-mTOR pathway (Figure S3A). Furthermore, *PTEN^mutant^* cells upregulated the expression of genes that correlate with radiation resistance, such as *SftpC, Slc34a2, Tub, Myh6* and *Shh,* while IR sensitizing genes, such as *Wisp2* and *Bex,* were enriched in KP tumors (Figure 3C). Additionally, *PTEN^mutant^* cells upregulate pathways associated with IR and Doxorubicin resistance (Figure 3D), both treatments resulting in double strand breaks. Genes associated with Telomere end packaging and maintenance were enriched in *PTEN^mutant^* cells compared to KP cells (Figure S3B). Overall, loss of *Pten* led to a transcriptomic shift towards pathways that are associated with aggressiveness, metastasis and therapy resistance (Figure 3D and S3A). This was further evidenced by increased expression of c-MYC (V1), E2F and reactive oxygen species (ROS) pathway genes in *KPP* tumor cells (Figure S3B).

**Figure 3:**
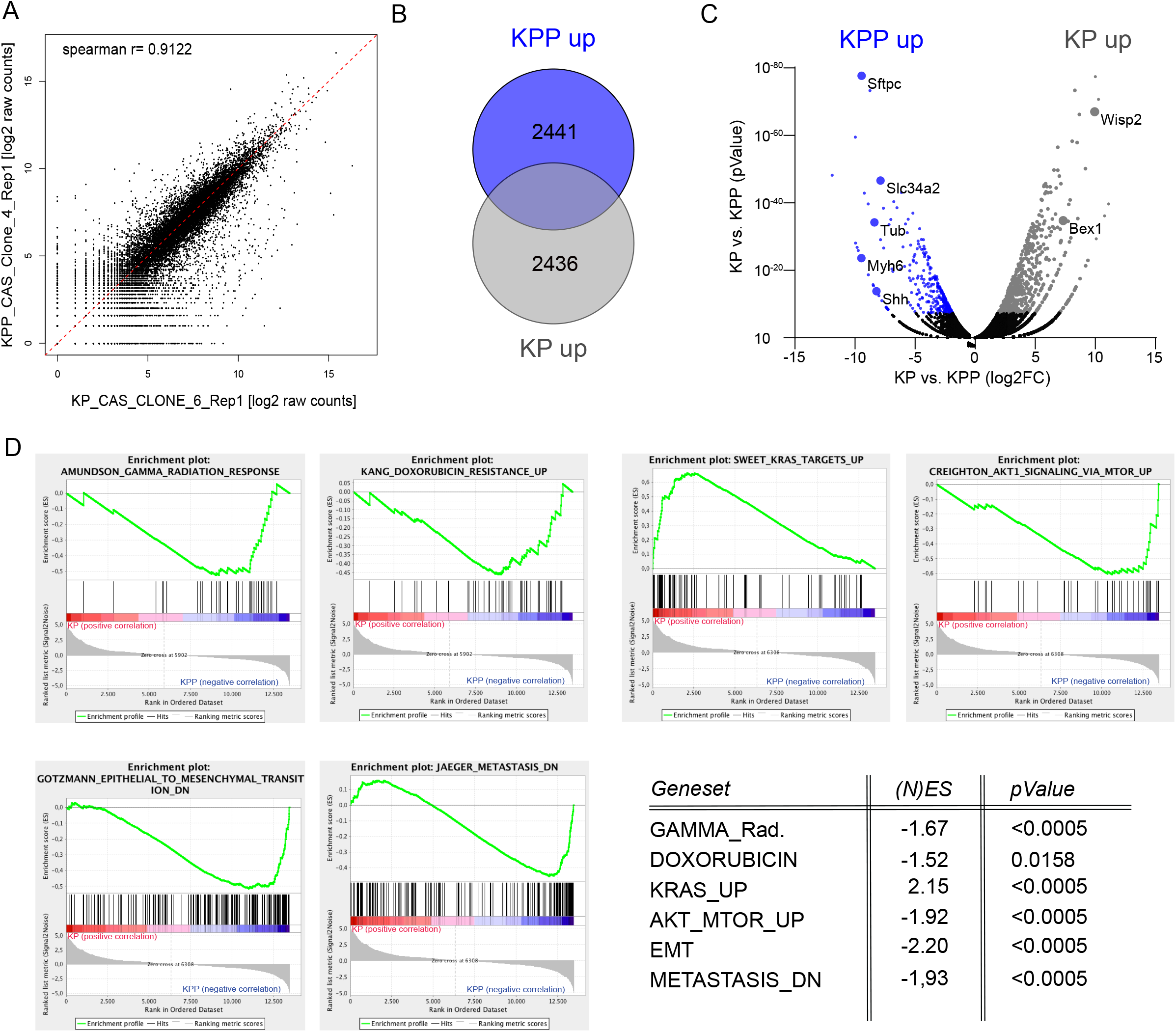
Loss of *Pten* alters DNA damage signalling pathways in murine NSCLC A) Correlation of gene expression changes of *KRas^G12D^:Trp53* (KP6) relative to *KRas^G12D^:Trp53:Pten* (KPP4). The diagonal line reflects a regression build on a linear model. R: Pearsons correlation coefficient. R=0.9122. B) Venn diagram of differentially up-regulated genes (log_2_FC>1.0 and q-value<0.05) between *KRas^G12D^:Trp53* (KP6) relative to *KRas^G12D^:Trp53:Pten* (KPP4). C) Volcano blot of differentially up- and downregulated genes in *KRas^G12D^:Trp53:Pten* (KPP4) relative to *KRas^G12D^:Trp53* (KP6).log_2_FCcut-off >1.0, −log10FC >1.5. Highlighted are genes involved in IR resistance; *SftpC, Slc34a2, Tub, Myh6* and *Shh,* or IR sensitivity, *Wisp2* and *Bex1*. n=3 D) Gene set enrichment analysis (GSEA) of Gamma radiation response, doxorubicin resistance up, KRas targets up, AKT1 signaling via mTOR, mesenchymal transition and metastasis *KRas^G12D^:Trp53* (KP) relative to *KRas^G12D^:Trp53:Pten* (KPP). n=3 each. Table with normalized enrichment score ((N)ES) and p-Value of GSEA. Also see Supplementary Figure S3.

*KPP* tumors appear to upregulate the DNA damage response already at steady state. To investigate if DNA damage recognition and clearance therefore varies between *Pten* proficient and deficient cells, we subjected *KP* and *KPP* cells to IR (8 Gy) and studied radiation induced presence and activity state of DNA damage kinase ATR as seen by phosphorylation, (Figure S3E). While non-irradiated cells had low amounts of phospho-ATR, already 5 minutes’ post IR exposure led to a significant and rapid increase of phospho-ATR in KP cells, while KPP failed to activate ATR (Figure S3C).

It is noteworthy that ATR is apparently not activated in *KPP* cells, while both cell lines upregulated γH2AX. This is an intriguing observation that could point towards an efficient mechanism for DNA damage recognition and clearance, present in Pten deficient tumor cells. This could putatively contribute to the DNA damage therapy evasion frequently observed in PTEN mutant patients. Furthermore, the lack of ATR activation during IR exposure argues that loss of PTEN could rewire the DNA damage signalling network towards DNA-PK or ATM.

### Interference with PI3K-mTORC signalling via the dual specific small molecule inhibitor PI-103 in PTEN^mutant^ cells

Loss of *PTEN* interferes with the PI3K–mTOR signalling cascade, leading to constant phosphorylation of AKT. Phospho-AKT activates DNA-PK, a key enzyme in DNA-damage recognition and repair^45, 46^. Cells may develop addiction to this situation. To investigate whether this could serve as an exploitable vulnerability, we irradiated the primary murine NSCLC cell lines KP and KPP in the presence or absence of PI-103, a potent PI3K/AKT and mTOR inhibitor, that also interferes with DNA-PK (Figure 4A and S4A, B and ^47^). Cells were pre-treated with 2 μM PI-103 for 3 h, followed by irradiation with 8 Gy. Whole protein extracts were collected at indicated time points post IR, followed by immunoblotting against total and phosphorylated AKT (Figure 4A and S4A). While whole protein levels as well as phosphorylated amounts of AKT were not altered in KP cells upon exposure to 8 Gy in presence or absence of PI-103, KPP cells showed pathway inhibition at time of irradiation and for at least two hours post irradiation, as seen by diminished phosphorylation of AKT (Figure 4A). However, the pathway was swiftly reactivated within 4 h post irradiation and AKT phosphorylation was fully restored (Figure 4A). This demonstrates that blockage of the PI3K-AKT pathway via PI-103 only effected a brief pathway inhibition in *Pten^mutant^* cells. Radiation dose dependent colony formation of KPP was not different in the presence of 2 μM PI-103, while KP showed mild sensitization. (Figure 4B and S4B). To investigate the differential responses of KP and KPP cells to ionizing irradiation in the presence or absence of PI-103 treatment, next, we measured cell survival by trypan blue staining with an automated cell counter. Here, in a dose dependent fashion, overall cell numbers were reduced when cells were exposed to PI-103 (Figure 4C). The small molecule inhibitor did not induce cell death at lower concentrations but synergized with ionizing radiation in the *Pten* wild type cancer cell line KP in higher concentrations (10 μM to 20 μM), as seen by a decrease in viable cells. The *Pten^mutant^* KPP cell line only demonstrated an initial growth disadvantage and a mild reduction in cell viability, however, tolerated higher concentrations of PI-103 in combination with IR than KP (Figure 4C).

**Figure 4:**
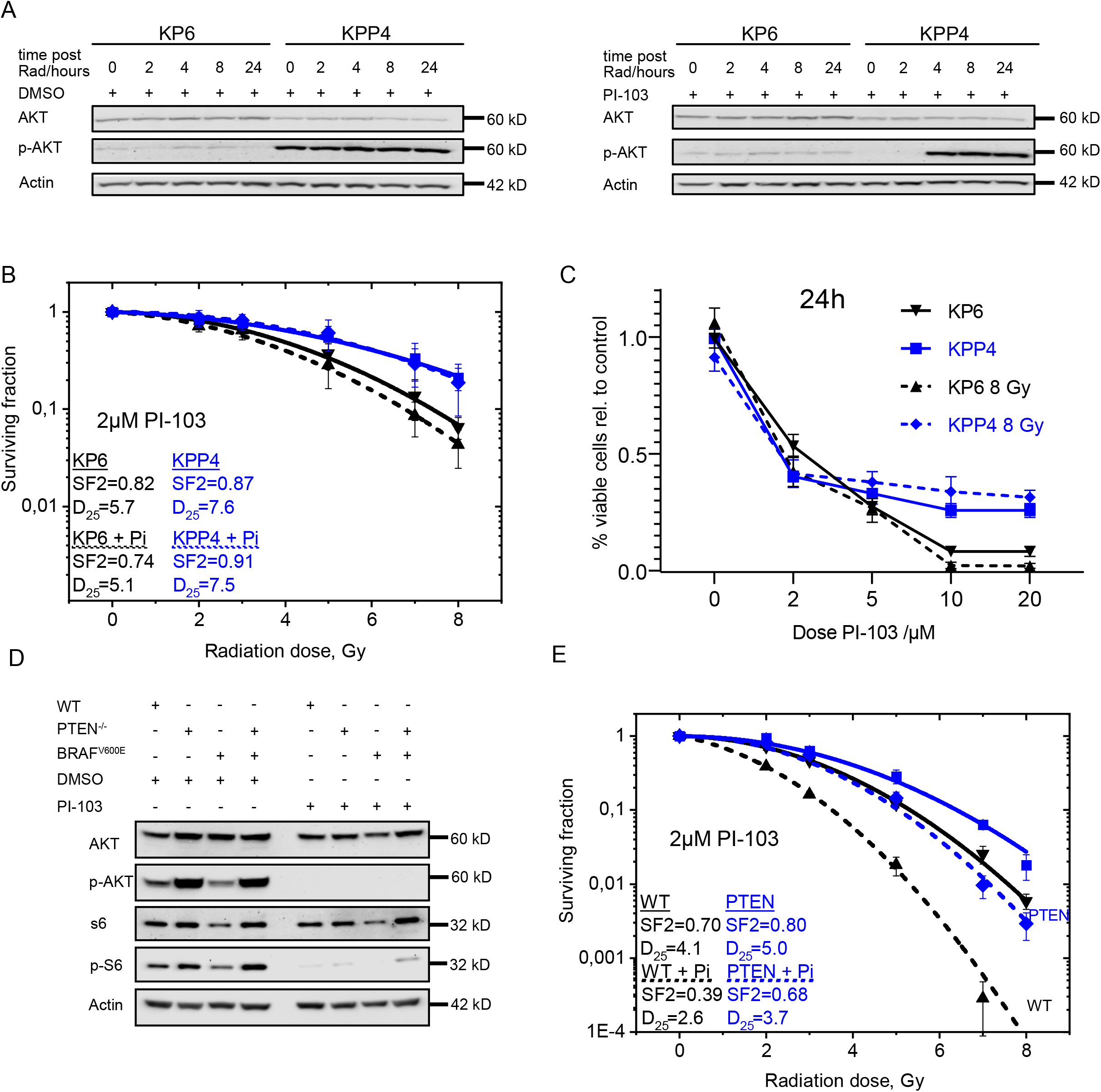
Impact of PI3K/mTOR inhibition in PTEN deficient cells A) Representative Immunoblot of KP6 and KPP4 cells without and with 2μM PI-103 treatment 3h before irradiation with 8 Gy at time points directly, 2h, 4h, 8h and 24h after irradiation. DMSO as solvent control. Actin and AKT as loading control. n=3. B) Colony formation assay KP6 (black) and KPP4 (blue) cells with 2 μM PI-103 (dashed lines) and DMSO as control (continuous lines) with re-seeding protocol (Figure S4A). SF 2: Surviving fraction at 2 Gy. D_25_: Dose in Gy with 25% survival. Error bars: Standard deviation. n=3. C) Relative number of living of KP6 (black) and KPP4 (blue) cells 27h after treatment with PI-103 in different concentrations, DMSO as control and 24 h after radiation with 8 Gy (dashed lines) or without radiation (continuous lines) (dead cells stained with trypan blue excluded from analysis). Error bars: Standard deviation. n=3. D) Immunoblot of (phospho-)AKT and (phospho-)S6 BEAS-2B wildtype (WT), *PTEN^homo^*, *BRAF^V600E^* and compound mutant cell lines without and with 2 μM PI-103 pre-treatment for 3 h. DMSO as solvent control. Actin serves as loading control. E) Colony formation assay of WT (black) and PTEN deficient (blue) BEAS-2B cells with 3 h pre-treatment of 2 μM PI-103 (dashed lines) and DMSO as control (continuous lines) with 24 h re-seeding protocol (Figure S4A). SF 2: Surviving fraction at 2 Gy. D_25_: Dose in Gy with 25% survival. Error bars: Standard deviation. n=3. Also see Supplementary Figure S4.

Treatment of BEAS-2B cells revealed slightly differing results. While solvent/DMSO treated cells showed robust activation of the AKT-mTORC pathway, as demonstrated by phosphorylation of AKT and S6 in *PTEN^mutant^* cells, exposure to 2 μM PI-103 for 3 h inhibited AKT and significantly reduced phosphorylation of S6 (Figure 4D). Dose dependent clonogenic survival upon IR in the presence of solvent control or PI-103 (Figure S4B) demonstrated, that treatment with PI-103 reduced IR resistance only to modest extent in PTEN deficient cells (both *PTEN^mutant^* and PTEN^mutant^ BRAF^V600E)^, while PTEN WT cells showed a distinct sensitisation to radiation (Figure 4E, S4C and D).

Our data demonstrate that combined PI3K, mTOR and DNA-PK inhibition is not an effective treatment to overcome *PTEN^mutant^* induced radiation resistance.

### Inhibition of ATM kinase by KU-60019 or AZD 1390 restores IR sensitivity in Pten^mut^ BEAS-2B and murine NSCLC cells

Next, we tested whether the DNA damage kinase ATM could present a target in *PTEN^mut^* cells. Two ATM inhibitors (KU-60019 and AZD 1390) were employed in our genetically engineered BEAS-2B and KP6 versus KPP4 cells (Figure 5A, B and S5A, B). Cells were treated with ATM inhibitor or solvent control for 27 hours (to model 3 hours pre-treatment and 24 hours of IR and recovery time), then dose dependent colony survival was measured. KU-60019 and AZD 1390 had little effect on overall cell survival up to a concentration of 3 μM in the tested cell lines, and growth inhibition was only observed in concentrations exceeding 10 μM (Figure 5A, B and S5A, B). Immunoblotting of genetically engineered BEAS-2B as well as KP versus KPP cells showed that non-irradiated cells had very low levels of detectable phosphorylated ATM or γH2AX (Figure 5C, D and S5C). Upon exposure to 8 Gy, phosphorylated ATM as well as γH2AX were strongly increased in all analysed cell lines. Treatment with 0.3 μM KU-60019 significantly reduced, and exposure to 3 μM KU-60019 blocked the phosphorylation of ATM and led to a marked reduction in overall γH2AX protein levels (Figure 5C, D). Loss of γH2AX indicates that interference with ATM activation in irradiated cells impairs downstream DNA damage signalling. Comparable results were obtained when AZD 1390 was used (Figure S5C). In concentrations exciding 3μM, AZD 1390 interfered with AKT phosphorylation in *Pten* mutant cells, potentially via blocking PI3K (Figure S5C).

**Figure 5:**
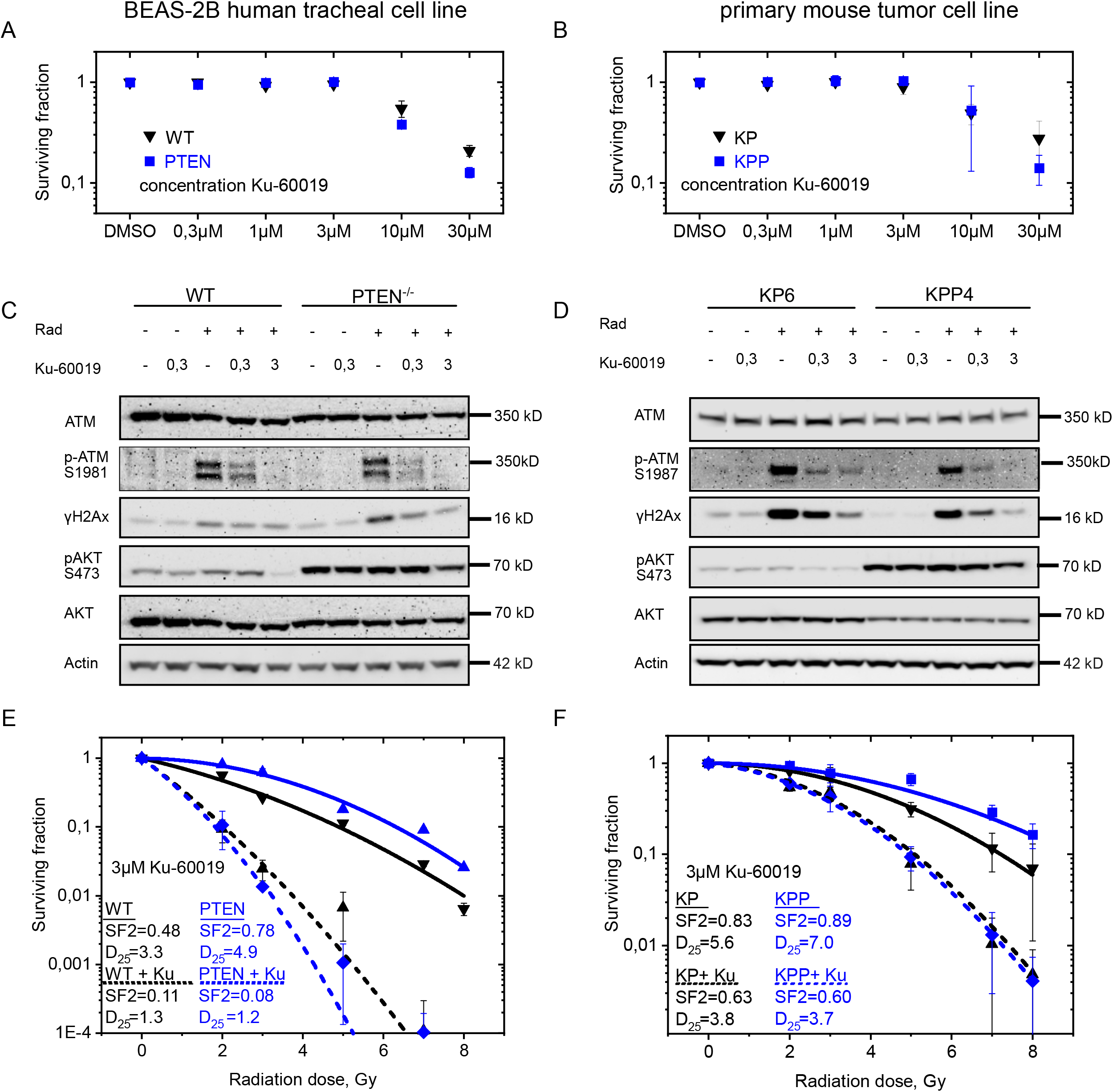
Impact of ATM inhibition in PTEN deficient cells A) Dose response of BEAS-2B WT (black) and BEAS-2B PTEN^*homo*^ (blue) BEAS-2B cells on colony forming ability following treatment with KU-60019 in different concentrations. Error bars: Standard deviation. n=3. B) Dose response of murine PTEN proficient KP6 (black) and PTEN deficient KPP4 cells on colony forming ability following treatment with KU-60019 in different concentrations. Error bars: Standard deviation. n=3. C) Immunoblot of WT and PTEN deficient BEAS-2B cells 30 min after irradiation with 8 Gy and 3 h pre-treatment with 0,3 μM and 3 μM KU-60019 before irradiation. DMSO as solvent control. Actin, ATM and AKT as loading control. n=3. D) Immunoblot of murine PTEN proficient KP6 and PTEN deficient KPP4 cells 30 min after irradiation with 8 Gy and 3 h pre-treatment with 0,3 μM and 3 μM KU-60019 before irradiation. DMSO as solvent control. Actin, ATM and AKT as loading control. n=3. E) Colony formation assay of WT (black) and PTEN deficient (blue) BEAS-2B cells with 3 h pre-treatment of 3 μM KU-60019 (dashed lines) and DMSO as control (continuous lines) with 24 h re-seeding protocol (Figure S4A). SF 2: Surviving fraction at 2 Gy. D_25_: Dose in Gy with 25% survival. Error bars: Standard deviation. n=3. F) Colony formation assay of murine PTEN proficient KP6 (black) and PTEN deficient KPP4 cells with 3 h pre-treatment 3 μM KU-60019 (dashed lines) and DMSO as control (continuous lines) with re-seeding protocol (Figure S4A). SF 2: Surviving fraction at 2 Gy. D_25_: Dose in Gy with 25% survival. Error bars: Standard deviation. n=3. Also see Supplementary Figure S5.

Next, we tested the combinatorial treatment of *PTEN/Pten* wild type and mutant cells with ATM inhibition and IR. To this end, cells were pre-treated with the indicated ATM inhibitors for 3 hours at nontoxic concentrations of 0.3 μM or 3 μM, respectively, followed by exposure to indicated doses of ionizing radiation. Cells were re-seeded and colony formation capacity was analysed. Exposure of *PTEN/Pten* mutant cells to ATM inhibitor, in an ATM-inhibitor dosage dependent fashion, resulted in radio-sensitization and reduction of clonogenic survival (Figure 5E, F and S5D, E). Comparable results were obtained when AZD 1390 was used (Figure S5F). It is note worth noting that the expression of oncogenic *BRAF^V600E^* did not alter the response of *PTEN* mutant cells to combinatorial treatment (Figure S5E).

These data demonstrate that ionizing radiation resistant *PTEN^mutant^* cells are addicted to the DNA damage kinase ATM. This tumor bottleneck can be exploited, as wild type nor tumor cells relied on active ATM for cell proliferation, at least *ex vivo*, and tolerated ATM inhibition via KU-60019 or AZD 1390, while in combination with ionizing radiation, *PTEN^mutant^* cells, human and murine, succumbed to therapy

### Pten^mut^ NSCLC require ATM to suppress a pro-apoptotic program upon IR

To gather further insights into how loss of *Pten/PTEN* reshapes the cellular responses upon ionizing radiation, we compared global changes in the appearance of phosphorylation, a major post-translational modification, required to regulate the activity of several key enzymes of the DNA damage response (DDR) and apoptosis signalling cascade^37, 48, 49^.

Analysis of the global phospho-proteome revealed fundamental differences between *Pten* proficient and deficient cell lines (Figure 6A and S6A). Already under basal conditions pathways associated with RNA splicing, apoptosis, RNA polymerase and stress responses were differentially regulated (Figure 6A and S6A). These steady-state differences might influence the reaction of these cells to stressors, such as radiation. Exposure to IR differentially regulated pathways associated with cell cycle and G2/M checkpoints, but also RNA Pol I & II, mRNA processing and TP53 activity or apoptosis (Figure 6B). Comparative phospho-proteomic analysis revealed a small cluster of apoptotic hallmark genes (MSigDB), differentially regulated after IR in *Pten* deficient cells (Figure 6C). Addition of ATM inhibitor KU-60019 resolved this deregulated cluster towards a *Pten* proficient like response (Figure 6C). Analysis of this cluster showed that pro-apoptotic proteins, such as Rara, Caspase 8, Diablo, Bax and Bcl2l1 were less phosphorylated in *Pten* deficient cells upon exposure to IR, hence, pro-apoptotic signalling was impaired (Figure 6D). Furthermore, KPP, when compared to KP, deregulated cell cycle checkpoint proteins, apoptosis, mRNA splicing and chromatid cohesion differentially to KP cells, thereby contributing to the increased tolerance towards ionizing radiation (Figure S6B). Addition of the small molecule ATM inhibitor KU-60019, reverted the ‘underrepresentation’ of phosphorylation of these factors and restored a pro-apoptotic signature in KPP to the same extend than KP (Figure 6D). The combination of IR and KU-60019 led to an increase in the phosphorylation of apoptotic execution phase proteins, apoptosis induced cleavage of proteins, cell cycle and death receptor signalling (Figure S6B).

**Figure 6:**
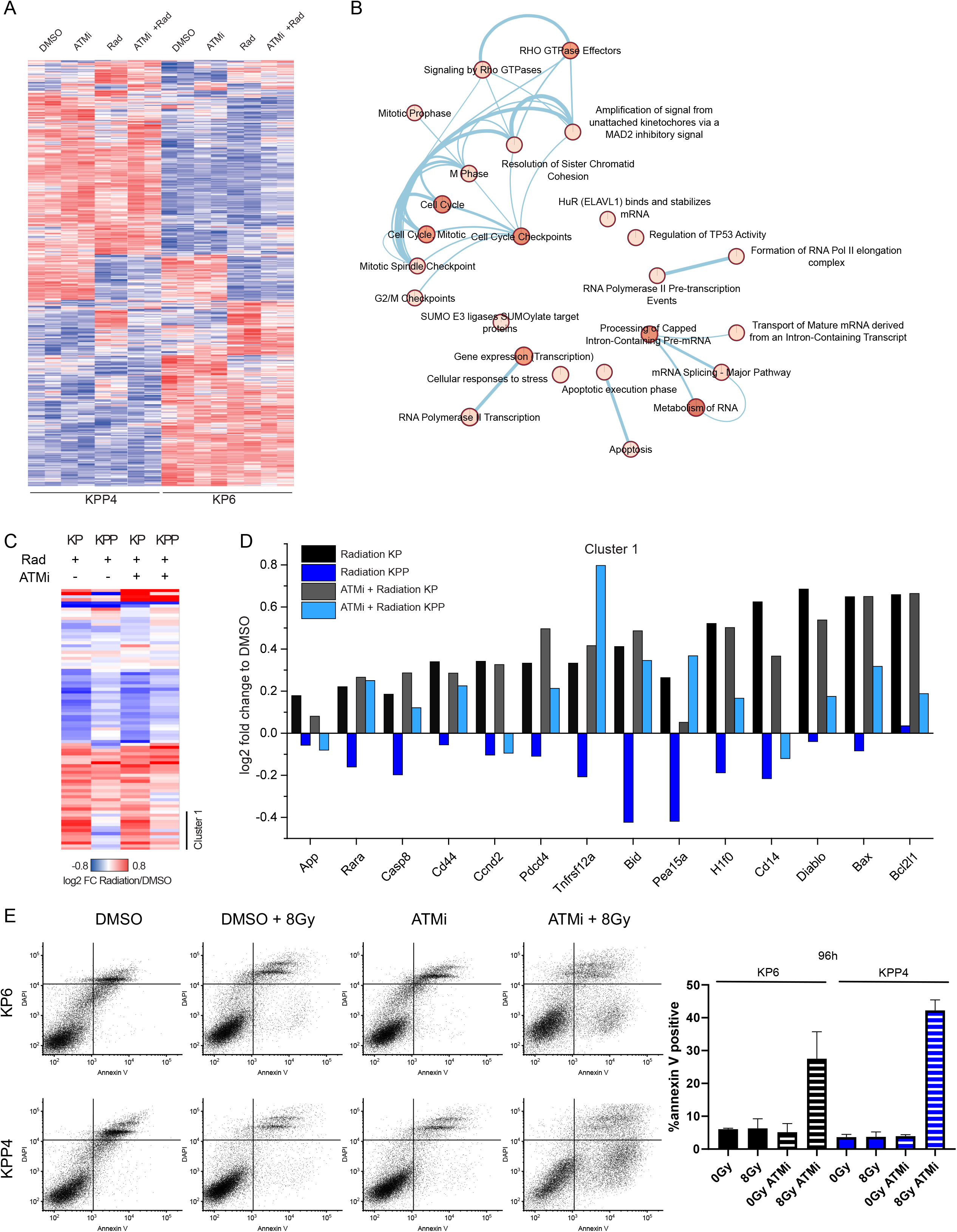
Multilevel proteomics show differential apoptosis signaling A) Heatmap of KP and KPP phosphorylation sites after treatment with solvent control, KU-60019, radiation and combined treatment. Phosphosites (rows) and samples (columns) have been hierarchically clustered using Euclidean distance. Quantification values have been standardized using Z-scoring to account for different scales. Color scales indicate Z-scores. B) Enrichment map showing Reactome pathways differentially regulated (log2 fold change differences >0.5) between KP and KPP cells upon radiation. Related pathways are connected by edges. Node coloring corresponds to ReactomeFI functional enrichment score. All pathways shown are significantly enriched with an FDR < 0.05. C) Heatmap showing total protein fold changes of apoptosis hallmark genes upon radiation and combinatorial treatment in KP and KPP cell lines. Clustering has been performed using hierarchical clustering with Euclidean distance. D) Bar graph showing log2 fold changes for genes identified in cluster I from C. The data indicates that combinatorial treatment rescues the expression differences upon radiation between the two cell lines. E) AnnexinV/DAPI staining of KP6 and KPP4 cells with 3 h pre-treatment 3 μM KU-60019 and DMSO as control with and without irradiation 8 Gy, 96h post irradiation. Supernatant of 96 h cultivation Medium was collected with trypsinized cells before staining. The lower right quadrant of the dot plots shows the apoptotic fraction measured with flow cytometer. The diagram shows the apoptotic fraction after 96h with different treatments. Error bars: Standard deviation. Also see Supplementary Figure S6.

To investigate if these effects indeed affect KP and KPP survival upon the combination of ionizing radiation and KU-60019, we performed fluorescent activated cell sorting (FACS) by using DAPI and the apoptosis marker Annexin V (Figure 6E). Exposure to 8 Gy ionizing radiation or the exposure to 3 μM KU-60019 had little effect on overall cell viability of KP or KPP cells (Figure 6E). Upon exposure to 8 Gy in combination with 3μM KU-60019, KP cells increased the percentage of cells in the apoptotic stage (to 30% Annexin V+/DAPI−, Figure 6E and S6C, D). KPP cells were more sensitive to the combinatorial treatment and showed a marked increase in apoptotic cells after 96 h, exceeding KP cells (>40% Annexin V+/DAPI−, Figure 6E and S6C, D).

These data demonstrate that ionizing radiation resistant *PTEN^mutant^* cells are addicted to the DNA damage kinase ATM, and ATR nor DNA-PK can substitute for ATM during therapy. This tumor bottleneck can be exploited, as wild type nor tumor cells relied on active ATM for cell proliferation, at least *ex vivo*, and tolerated ATM inhibition via KU-60019 or AZD 1390, while in combination with ionizing radiation *PTEN^mutant^* cells, human and murine, succumbed to therapy.

### Combining ionizing radiation with ATM inhibition results in PTEN^mutant^ tumor regression in ex vivo organotypic lung tumor slice cultures

The *in vitro* result was reproduced in a multicellular *ex vivo* organotypic lung system (Figure 7A and Figure S7A). Isogenic murine KP6 and KPP4 cells were orthotopically re-transplanted in immune-competent C57BL6/J mice (Figure 7A). 8 weeks post-transplantation, mice were sacrificed, and lungs analysed for tumor engraftment of green fluorescent protein positive (GFP^+^) tumor cells, followed by live tissue sectioning with a Leica V1200S vibratome and subsequent culture of life tissue sections in a 24 well plate (Figure 7A). Slices containing tumor (GFP^+^) and wild type tissue (GFP^−^) were cultured in standard medium (DMEM, 10 % FCS) and exposed to either IR (8 Gy), 3 μM KU-60019, or a combination of both, according to treatment regime, followed by imaging of GFP^+^ cells for indicated time points (Figure 7A and S7A and B). We used the GFP signal of the transplanted tumor cells as a surrogate marker for cell viability, as dead cells lose GFP signal.

**Figure 7:**
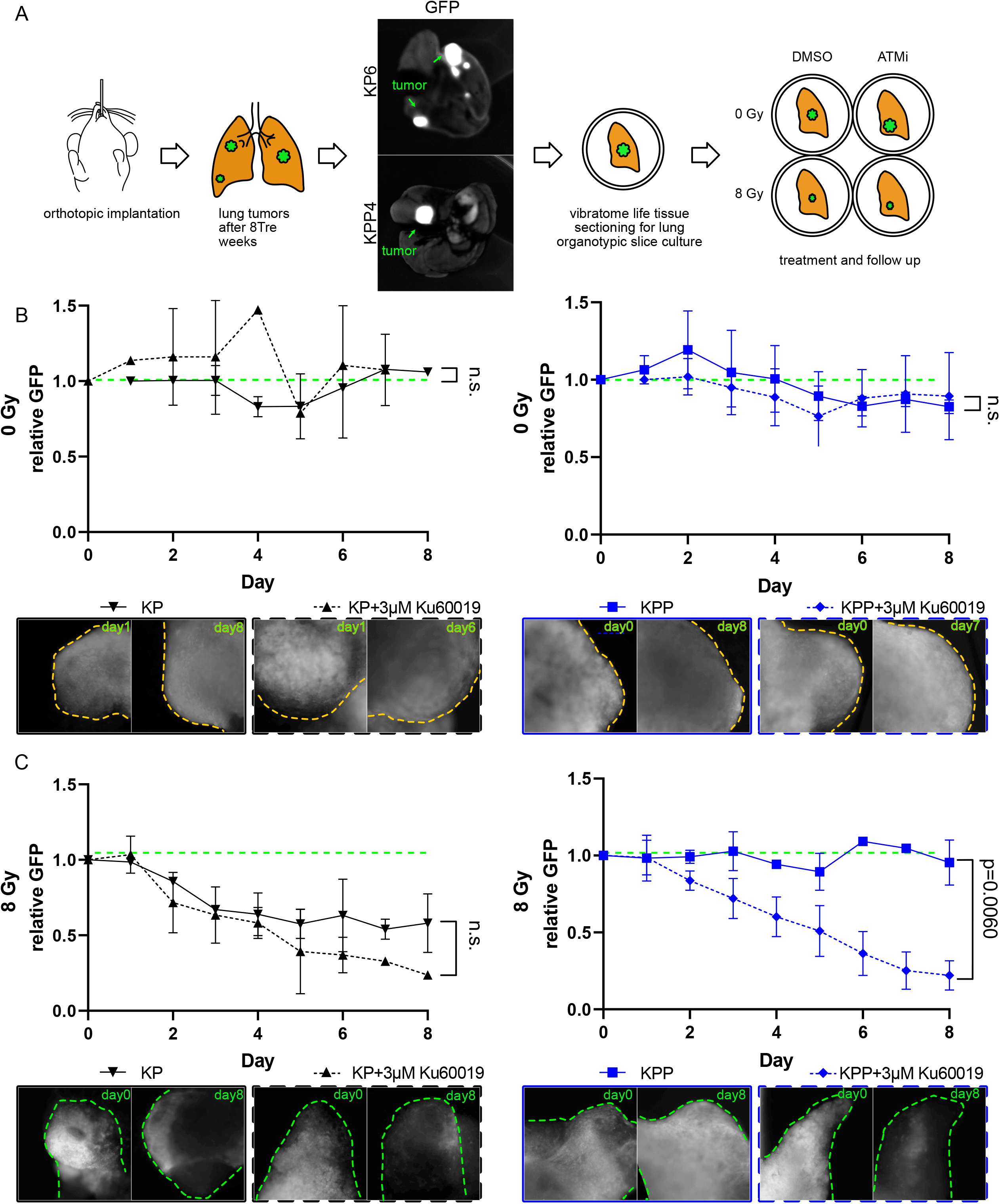
Tumor slice culture response to KU-60019 treatment and radiation A) Schematic of orthotopic transplantation of GFP positive KP6 and KPP4 cells to immune competent C57BL/6 mice. The picture shows GFP positive tumors in mouse lungs after 8 weeks. The tumor bearing mouse lungs were cut by vibratome and cultured in 10% FCS/DMEM in 24 well plates. Culture slices were treated with DMS0 or ATM inhibitor and irradiated with 0 Gy or 8 Gy. B) Tissue slices (n=2-4) of transplanted KP6 (black) and KPP4 (blue) were pre-treated with DMSO (continuous line) or 3 μM Ku60019 (dashed line) Treated tissue slices were observed and pictures of same tumor sites were taken for 8 days. The fluorescent signal of the tumor area was measured, and background area was subtracted. On the Graphs the GFP signal over time with different treatment conditions is shown. Below each graph are typical pictures of measured tumor sites illuminated with standardized 488 nm led light source and same camera settings. C) Tissue slices (n=2-4) of transplanted KP6 (black) and KPP4 (blue) were pre-treated with DMSO (continuous line) or 3 μM Ku60019 (dashed line) and irradiated with 8 Gy. Treated tissue slices were observed and pictures of same tumor sites were taken for 8 days. The fluorescent signal of the tumor area was measured, and background area was subtracted. On the Graphs the GFP signal over time with different treatment conditions is shown. Below each graph are typical pictures of measured tumor sites illuminated with standardized 488 nm led light source and same camera settings. Also see Supplementary Figure S7.

Culture of organotypic slices for 8 days showed no deterioration of the GFP signal of untreated KP and KPP tumor cells (Figure 7B and S7B). Exposure to 3 μM KU-60019 alone did not result in tumor cell death, as seen by consistent GFP intensity over the course of the experiment (Figure 7B). Exposure of KP transplant tumors to a single dose of IR (8 Gy) resulted in a reduction in overall GFP signal intensity, indicating that tumor cells died upon treatment, (Figure 7C). This effect was further enhanced by combining IR with 3 μM KU-60019 (Figure 7C and S7B). Exposure of KPP transplant tumors to IR alone showed no regression of GFP signal intensity, thereby confirming the therapy resistance of *PTEN^mutant^* cells in a multicellular system (Figure 7C and S7B). Combined treatment with 8 Gy and 3 μM KU-60019 led to a rapid decrease of the GFP signal in the *PTEN^mutant^* KPP tumors, that rapidly diminished comparable to *PTEN^wt^* (Figure 7C and S7B).

These data show that ATM inhibition potentiated IR therapy responses in tumor cells and re-establishes a sensitivity of otherwise radiation resistant *PTEN^mutant^* tumor cells. highlighting that targeting ATM could result in a general improvement of IR-based therapy.

## Discussion

Radiotherapy is an important modality in cancer treatment. Ionizing radiation inflicts DNA damage and challenges the complex DNA damage repair machinery in cells. Current knowledge identifies a complex network of more than 800 genes involved in damage recognition and handling, related cell cycle response and eventually removal of critically damaged cells. This network is characterized by redundancy and alternative and fallback pathways (e.g. repair of double strand breaks). From an evolutionary point of view this is of importance to maintain genomic stability and control of proliferation in multicellular organisms.

Tumor cells, in contrast to non-transformed cells, frequently harbor mutations in check point genes and fail to halt the cell cycle to initiate the repair of damaged DNA^50–53^. Mutations in oncogenes, such as *KRAS^54^*; and subsets of loss-of-function mutations in tumor suppressors, such as *FBXW7^44, 55^ or STK11^56^*, can cause resistance to DNA damage based therapies. Identification of exploitable ‘bottlenecks’ for tumor cell survival might be an option to advance our current treatment options.

One such exploitable bottleneck is presented by mutations in the phosphatase and tensin homologue (*PTEN*). Our analysis of publicly available datasets revealed that *PTEN* is frequently mutated in NSCLC, ranging from transcriptional downregulation to genetic loss, and frequently co-occur with gain of function mutations in the oncogene *KRAS* and loss of function mutations in the tumor suppressor *TP53*. *PTEN* gene dosage is a direct prognostic marker for therapy outcome and patient survival, as already a reduction in gene expression negatively correlated with patient survival and ionizing radiation therapy success for both NSCLC entities, adenocarcinoma and squamous cell carcinoma. This effect is not limited to NSCLC, but was reported in other tumor entities where *PTEN* was mutated e.g. glioblastoma and prostate tumors^57^. Genetic loss accelerates tumor growth, enhances tumor burden and shortens overall survival^58^.

Several murine *in vivo* systems were established to analyse the role of Pten in cancer progression and survival, such as pancreas^59, 60^, breast^61^, endometrium^62^ and lung^63^. We have established a novel mouse model using CRISPR gene editing to delete common tumor suppressors, such as *Trp53* and *Pten*, together with mutating *KRas* to *Kras^G12D^*, to establish primary tumors and cell lines. In this model, we reproduced both accelerated tumor growth and reduced survival caused after loss of *Pten*. This genetic alteration was sufficient also to enhance resistance towards ionizing radiation.

Analysis of *PTEN* dependent changes of the transcriptome in our primary murine lung tumor cells revealed that loss of Pten had a significant impact on gene expression. Gene sets associated with epithelial–mesenchymal transition (EMT) and metastasis were enriched together with increased expression of Myc target gene signatures as well as G2M checkpoint genes and E2F pathway members. Depletion of Pten also altered the expression of gene sets associated with therapy response, in particular against ionizing irradiation or doxorubicin treatment, both treatments causing double strand breaks and ROS^64, 65^. Loss of PTEN obviously profoundly changes the cellular environment if DNA damage is encountered.

Although handling of DNA-damage occurs in a complex and pleiotropic network, selective gene editing of PTEN using CRISPR/Cas technology led to modification of radiation sensitivity for a multitude of endpoints (clonogenic survival, cell number and apoptosis in cell culture and cell viability and tumor size in organ culture). The effect was found both in a presumable stable “normal” cell line (BEAS-2B) and in a “tumor” cell line harbouring additional mutations (KP; e.g. p53 and KRas). The specificity of the intervention was confirmed by reconstitution of PTEN function in the mutated clones via lentiviral transduction. A reversal of the effects on radiosensitivity was demonstrated.

In BEAS-2B cells, CRISPR gene editing reduced radiation sensitivity in *PTEN^hetzerozygous^* and in *PTEN^homozygous^* deficient cells. Our data suggest that IR resistance strongly correlates with PTEN gene and protein status. Our work also demonstrated that *PTEN* loss alone is sufficient to drive IR resistance, as the *in cellulo* gene modification in BEAS-2B allowed us to not only create *PTEN^hetzerozygous^* and *PTEN^homozygous^* mutant cells, but also to combine it with oncogenic drivers, such as *BRAF^V600E^*. In our experiments the overexpression of *BRAF^V600E^* had no effect on IR resistance in BEAS-2B wild type or *PTEN^mut^* single and compound cells, showing that MAPK pathway alteration has only low impact on radiation sensitivity in this cell system.

Loss of PTEN causes hyper-activation of pAKT and its downstream signals^66^. pAKT, apart from many other effects, activates DNA-PK, an important protein in the DNA damage repair cascade, especially in classical non homologous end joining (NHEJ). We used PI-103 to inhibit the PI3K pathway. However, PTEN^mut^ cells still showed lower sensitization to radiation treatment than their PTEN^wt^ counterpart. This was potentially due to a fast rebound of pAKT. Alternatively, backup pathways regulating DNA damage repair might be preferentially active in PTEN^mut^ cells.

Non-transformed and oncogenic transformed cells rely on an efficient mechanism to identify and repair damaged DNA. The major DNA kinases, DNA-PK^66^, ATR^67^ and ATM^68^, recognize various types of damage, ranging from interstrand crosslink to single- and double stand breaks, and initiate downstream repair pathways, such as non-homologous end joining or homologous repair^69^.

In mammalian cells NHEJ is the dominant way of repairing DNA-double strand breaks. This constitutes a fast, partly error prone mechanism. Recent data show that fidelity and effectiveness of NHEJ depends on the extent of microhomology and overlaps with a further DNA-PK independent repair pathway (alternative-end joining). Interestingly, inhibition of DNA-PK via the compound PI-103 had no effect on IR resistance of KP nor KPP cells and only marginally induced IR sensitivity in BEAS-2B single and compound mutant cells.

So called toxic non-homologous end joining has been identified in ATM-deficient models if ATR was inhibited. We discuss a similar scenario in our models, where in PTEN deficient cells IR failed to elicit ATR activation as backup repair pathway, and subsequent ATM-inhibition caused strong radio sensitization.

The failure to activate ATR upon exposure to ionizing radiation was unexpected. However, PTEN is a key signal transducer and has functions independent of its proliferation directed cytoplasmic lipid phosphatase activity. Recent studies suggest that PTEN also localizes in the nucleus and is involved in chromatin functions^70^. Ma et al. showed that phosphorylation of PTEN at tyrosine 240 enhanced DDR via Rad51 and homologous end joining repair^71^.

Hence, we assume that KPP and BEAS-2B PTEN^*mut*^ with loss of PTEN function, mainly relied on ATM and NHEJ to sense and resolve DNA damage after irradiation. Treatment with ATM-inhibitors KU-60019 and AZD 1390 in low concentrations had no effect on cell survival or proliferation of Pten mutant and wild type cells. However, in combination with ionizing radiation it enhanced radiation sensitivity disproportionately in PTEN^*mut*^ and abolished the difference to the wild type. This effect was demonstrated both in classical cell culture and in an organotypic multicellular system. The combined treatment of PTEN^*mut*^ NSCLC by IR and ATM inhibition led to marked tumor regression.

This combination is synergistic and seems especially active in PTEN deficient tumors. While ATM inhibitors can be given with low systemic side effects, modern radiotherapy localizes treatment to the tumor with tight margins. This could create “spatial cooperation” in otherwise relatively radiation resistant tumors. A first clinical trial is evaluating tolerance of ATM inhibitors and radiation therapy (NCT03423628). The trial does not stratify treatment for different genetic backgrounds and therefore could miss significant improvement for selected but common groups, like patients with PTEN deficient tumors. We suggest that genetic stratification and personalized treatment might gain importance also in radiation therapy. PTEN and ATM are already part of clinically established tumor sequencing panels and results should find access to therapeutic decisions.

## Conclusion

In this study, we investigated the role of PTEN in response to radiation induced damage by genetically modulating PTEN in the human tracheal stem cell like cell line BEAS-2B. We observed in compound mutant cell lines that the IR resistance phenotype of PTEN-deficient tumors is indeed dictated by alterations in PTEN alone. This was validated in murine models of NSCLC, where loss of Pten induced IR resistance as well. The effect was not resolved by inhibition of DNA-PK and independent of ATR activation. However, pharmacological ATM inhibition (via the small molecules KU-60019 or AZD 1390) was able to increase radiation sensitivity and pointed to a crucial role of the DNA damage kinase ATM in a PTEN-deficient situation. These results from monolayer cell culture were reproduced *in* and *ex vivo* organoptypic slice culture assay. Analysis of transcriptional changes upon PTEN loss and obvious differences in activation of γH2AX points to shifts in DNA damage detection and response and resulting synthetic lethality in PTEN-deficient tumors. Our study suggests that tumors harbouring a loss of function mutation in PTEN can be therapeutically addressed by irradiation in combination with ATM inhibition.

## Supporting information

STAR Methods

## abbreviations

DMSO: dimethyl sulfoxide
RER: radiation enhancement ratio
DER: dose enhancement ratio
KP: *KRas^G12D^ :Tp53^mut^*
KPP: *KRas^G12D^ :Tp53^mut^:Pten^mut^*
H&E: haematoxylin and eosin
NSCLC: non-small cell lung cancer
UICC: union internationale contre le cancer
NGS: next generation sequencing
PTEN: phosphatase and tensin homologue
ROS: reactive oxygen species
UICC: union internationale contre le cancer
IR: ionizing radiation
ATM: ataxia telangiectasia mutated kinase
ADC: adenocarcinoma
SCC: squamous cell carcinoma
TMB: tumor mutational burden
AAV: adeno-associated virus
GSEA: gene set enrichment analysis
DDR: DNA damage response
FACS: fluorescent activated cell sorting
GFP: green fluorescent protein
NHEJ: homologous end joining

## Acknowledgements

We are grateful to the animal facility and Barbara Bauer at the Biocenter, University Würzburg. We thank Clare C. Davies from University of Birmingham for critical suggestions and discussions. C.P.G. and O.H. are supported by the German Cancer Aid via grant 70112491, M.R. is funded by the DFG-GRK 2243 and IZKF B335. M.E.D. and M.R. are funded by the German Israeli Foundation grant 1431. T. F. is funded by the IZKF program Z2/CS-1.

## Author contributions

Conceptualization: T.F., M.E.D.; Methodology: T.F. (in vitro) and O.H. (*in vivo*), C.S.V. (Operetta system); Formal analysis: C.P.G. and M.Re. (Bioinformatics), M.Ro. and M.E.D. (Pathology); Investigation: T.F., O.H., M.Re., C.P.G. B.P. M.Ro., M.E.D. Resources: M.Ro., M.F., M.E.D.; Writing-original draft: M.E.D.; Writing-review and editing: T.F., O.H., M.Re., M.Ro, M.F., M.E.D.; Supervision: M.E.D.; Funding acquisition: T.F., M.F., M.E.D.

## REFERENCES

1. Kohlbrenner, E. et al. Quantification of AAV particle titers by infrared fluorescence scanning of coomassie-stained sodium dodecyl sulfate-polyacrylamide gels. Hum Gene Ther Methods 23, 198–203 (2012).

2. Buchel, G. et al. Association with Aurora-A Controls N-MYC-Dependent Promoter Escape and Pause Release of RNA Polymerase II during the Cell Cycle. Cell Rep 21, 3483–3497 (2017).

3. Kim, D. & Salzberg, S.L. TopHat-Fusion: an algorithm for discovery of novel fusion transcripts. Genome Biol 12, R72 (2011).

4. Langdon, W.B. Performance of genetic programming optimised Bowtie2 on genome comparison and analytic testing (GCAT) benchmarks. BioData Min 8, 1 (2015).

5. Robinson, M.D., McCarthy, D.J. & Smyth, G.K. edgeR: a Bioconductor package for differential expression analysis of digital gene expression data. Bioinformatics 26, 139–140 (2010).

6. Mi, H., Muruganujan, A. & Thomas, P.D. PANTHER in 2013: modeling the evolution of gene function, and other gene attributes, in the context of phylogenetic trees. Nucleic Acids Res 41, D377–386 (2013).

7. Gao, J. et al. Integrative analysis of complex cancer genomics and clinical profiles using the cBioPortal. Sci Signal 6, pl1 (2013).

8. Cerami, E. et al. The cBio cancer genomics portal: an open platform for exploring multidimensional cancer genomics data. Cancer Discov 2, 401–404 (2012).

9. Nagy, A., Munkacsy, G. & Gyorffy, B. Pancancer survival analysis of cancer hallmark genes. Sci Rep 11, 6047 (2021).

10. McAlister, G.C. et al. MultiNotch MS3 enables accurate, sensitive, and multiplexed detection of differential expression across cancer cell line proteomes. Anal Chem 86, 7150–7158 (2014).

11. Ferlay, J. et al. Estimating the global cancer incidence and mortality in 2018: GLOBOCAN sources and methods. Int J Cancer 144, 1941–1953 (2019).

12. Cancer Genome Atlas Research, N. et al. The Cancer Genome Atlas Pan-Cancer analysis project. Nat Genet 45, 1113–1120 (2013).

13. Cancer Genome Atlas Research, N. Comprehensive molecular profiling of lung adenocarcinoma. Nature 511, 543–550 (2014).

14. Cancer Genome Atlas Research, N. Comprehensive genomic characterization of squamous cell lung cancers. Nature 489, 519–525 (2012).

15. Moreira, A.L. & Eng, J. Personalized therapy for lung cancer. Chest 146, 1649–1657 (2014).

16. Pakkala, S. & Ramalingam, S.S. Personalized therapy for lung cancer: striking a moving target. JCI Insight 3 (2018).

17. McDonald, F. et al. Management of stage I and II nonsmall cell lung cancer. Eur Respir J 49 (2017).

18. Siegel, R.L., Miller, K.D. & Jemal, A. Cancer statistics, 2018. CA Cancer J Clin 68, 7–30 (2018).

19. Varlotto, J.M. et al. Failure rates and patterns of recurrence in patients with resected N1 non-small-cell lung cancer. Int J Radiat Oncol Biol Phys 81, 353–359 (2011).

20. Taugner, J. et al. Pattern-of-failure and salvage treatment analysis after chemoradiotherapy for inoperable stage III non-small cell lung cancer. Radiat Oncol 15, 148 (2020).

21. Gajra, A. et al. Time-to-Treatment-Failure and Related Outcomes Among 1000+ Advanced Non-Small Cell Lung Cancer Patients: Comparisons Between Older Versus Younger Patients (Alliance A151711). J Thorac Oncol 13, 996–1003 (2018).

22. Liu, L. et al. PTEN inhibits non-small cell lung cancer cell growth by promoting G0/G1 arrest and cell apoptosis. Oncol Lett 17, 1333–1340 (2019).

23. Hamarsheh, S., Gross, O., Brummer, T. & Zeiser, R. Immune modulatory effects of oncogenic KRAS in cancer. Nat Commun 11, 5439 (2020).

24. Papillon-Cavanagh, S., Doshi, P., Dobrin, R., Szustakowski, J. & Walsh, A.M. STK11 and KEAP1 mutations as prognostic biomarkers in an observational real-world lung adenocarcinoma cohort. ESMO Open 5 (2020).

25. Chen, C.Y., Chen, J., He, L. & Stiles, B.L. PTEN: Tumor Suppressor and Metabolic Regulator. Front Endocrinol (Lausanne) 9, 338 (2018).

26. Lee, Y.R., Chen, M. & Pandolfi, P.P. The functions and regulation of the PTEN tumour suppressor: new modes and prospects. Nat Rev Mol Cell Biol 19, 547–562 (2018).

27. Xiao, J. et al. PTEN expression is a prognostic marker for patients with non-small cell lung cancer: a systematic review and meta-analysis of the literature. Oncotarget 7, 57832–57840 (2016).

28. Chang, L. et al. PI3K/Akt/mTOR pathway inhibitors enhance radiosensitivity in radioresistant prostate cancer cells through inducing apoptosis, reducing autophagy, suppressing NHEJ and HR repair pathways. Cell Death Dis 5, e1437 (2014).

29. Vidotto, T. et al. Emerging role of PTEN loss in evasion of the immune response to tumours. Br J Cancer 122, 1732–1743 (2020).

30. Vivanco, I. et al. The phosphatase and tensin homolog regulates epidermal growth factor receptor (EGFR) inhibitor response by targeting EGFR for degradation. Proc Natl Acad Sci U S A 107, 6459–6464 (2010).

31. Hou, S.Q., Ouyang, M., Brandmaier, A., Hao, H. & Shen, W.H. PTEN in the maintenance of genome integrity: From DNA replication to chromosome segregation. Bioessays 39 (2017).

32. Song, M.S. et al. Nuclear PTEN regulates the APC-CDH1 tumor-suppressive complex in a phosphatase-independent manner. Cell 144, 187–199 (2011).

33. Chen, Z.H. et al. PTEN interacts with histone H1 and controls chromatin condensation. Cell Rep 8, 2003–2014 (2014).

34. Sun, Z. et al. PTEN C-terminal deletion causes genomic instability and tumor development. Cell Rep 6, 844–854 (2014).

35. Furdui, C.M. Ionizing radiation: mechanisms and therapeutics. Antioxid Redox Signal 21, 218–220 (2014).

36. Borrego-Soto, G., Ortiz-Lopez, R. & Rojas-Martinez, A. Ionizing radiation-induced DNA injury and damage detection in patients with breast cancer. Genet Mol Biol 38, 420–432 (2015).

37. Huang, R.X. & Zhou, P.K. DNA damage response signaling pathways and targets for radiotherapy sensitization in cancer. Signal Transduct Target Ther 5, 60 (2020).

38. Canman, C.E. et al. Activation of the ATM kinase by ionizing radiation and phosphorylation of p53. Science 281, 1677–1679 (1998).

39. Tribius, S., Pidel, A. & Casper, D. ATM protein expression correlates with radioresistance in primary glioblastoma cells in culture. Int J Radiat Oncol Biol Phys 50, 511–523 (2001).

40. Ito, K. et al. Regulation of reactive oxygen species by Atm is essential for proper response to DNA double-strand breaks in lymphocytes. J Immunol 178, 103–110 (2007).

41. Li, K. et al. ATM inhibition induces synthetic lethality and enhances sensitivity of PTEN-deficient breast cancer cells to cisplatin. Exp Cell Res 366, 24–33 (2018).

42. Chen, J.H. et al. ATM-mediated PTEN phosphorylation promotes PTEN nuclear translocation and autophagy in response to DNA-damaging agents in cancer cells. Autophagy 11, 239–252 (2015).

43. Hartmann, O. et al. Implementation of CRISPR/Cas9 Genome Editing to Generate Murine Lung Cancer Models That Depict the Mutational Landscape of Human Disease. Front Cell Dev Biol 9, 641618 (2021).

44. Prieto-Garcia, C. et al. Maintaining protein stability of Np63 via USP28 is required by squamous cancer cells. EMBO Mol Med 12, e11101 (2020).

45. Brunner, A. et al. PTEN and DNA-PK determine sensitivity and recovery in response to WEE1 inhibition in human breast cancer. Elife 9 (2020).

46. Szymonowicz, K., Oeck, S., Malewicz, N.M. & Jendrossek, V. New Insights into Protein Kinase B/Akt Signaling: Role of Localized Akt Activation and Compartment-Specific Target Proteins for the Cellular Radiation Response. Cancers (Basel) 10 (2018).

47. Park, S. et al. PI-103, a dual inhibitor of Class IA phosphatidylinositide 3-kinase and mTOR, has antileukemic activity in AML. Leukemia 22, 1698–1706 (2008).

48. Brinkmann, K., Schell, M., Hoppe, T. & Kashkar, H. Regulation of the DNA damage response by ubiquitin conjugation. Front Genet 6, 98 (2015).

49. Marechal, A. & Zou, L. DNA damage sensing by the ATM and ATR kinases. Cold Spring Harb Perspect Biol 5 (2013).

50. Bhattacharya, S. & Asaithamby, A. Repurposing DNA repair factors to eradicate tumor cells upon radiotherapy. Transl Cancer Res 6, S822–S839 (2017).

51. Medema, R.H. & Macurek, L. Checkpoint control and cancer. Oncogene 31, 2601–2613 (2012).

52. Nikolaev, A. & Yang, E.S. The Impact of DNA Repair Pathways in Cancer Biology and Therapy. Cancers (Basel) 9 (2017).

53. Khanna, A. DNA damage in cancer therapeutics: a boon or a curse? Cancer Res 75, 2133–2138 (2015).

54. Wang, M. et al. Radiation Resistance in KRAS-Mutated Lung Cancer Is Enabled by Stem-like Properties Mediated by an Osteopontin-EGFR Pathway. Cancer Res 77, 2018–2028 (2017).

55. Ruiz, E.J. et al. LUBAC determines chemotherapy resistance in squamous cell lung cancer. J Exp Med 216, 450–465 (2019).

56. Sitthideatphaiboon, P. et al. LKB1 mutations in NSCLC are associated with KEAP1/NRF2-dependent radiotherapy resistance targetable by glutaminase inhibition. Clin Cancer Res (2020).

57. McCabe, N. et al. Mechanistic Rationale to Target PTEN-Deficient Tumor Cells with Inhibitors of the DNA Damage Response Kinase ATM. Cancer Res 75, 2159–2165 (2015).

58. Bazzichetto, C. et al. PTEN as a Prognostic/Predictive Biomarker in Cancer: An Unfulfilled Promise? Cancers (Basel) 11 (2019).

59. Hill, R. et al. PTEN loss accelerates KrasG12D-induced pancreatic cancer development. Cancer Res 70, 7114–7124 (2010).

60. Rosenfeldt, M.T. et al. PTEN deficiency permits the formation of pancreatic cancer in the absence of autophagy. Cell Death Differ 24, 1303–1304 (2017).

61. Ebbesen, S.H. et al. Pten loss promotes MAPK pathway dependency in HER2/neu breast carcinomas. Proc Natl Acad Sci U S A 113, 3030–3035 (2016).

62. Cheng, H. et al. A genetic mouse model of invasive endometrial cancer driven by concurrent loss of Pten and Lkb1 Is highly responsive to mTOR inhibition. Cancer Res 74, 15–23 (2014).

63. Iwanaga, K. et al. Pten inactivation accelerates oncogenic K-ras-initiated tumorigenesis in a mouse model of lung cancer. Cancer Res 68, 1119–1127 (2008).

64. Kim, S.Y. et al. Doxorubicin-induced reactive oxygen species generation and intracellular Ca2+ increase are reciprocally modulated in rat cardiomyocytes. Exp Mol Med 38, 535–545 (2006).

65. Perillo, B. et al. ROS in cancer therapy: the bright side of the moon. Exp Mol Med 52, 192–203 (2020).

66. Mohiuddin, I.S. & Kang, M.H. DNA-PK as an Emerging Therapeutic Target in Cancer. Front Oncol 9, 635 (2019).

67. Kim, D., Liu, Y., Oberly, S., Freire, R. & Smolka, M.B. ATR-mediated proteome remodeling is a major determinant of homologous recombination capacity in cancer cells. Nucleic Acids Res 46, 8311–8325 (2018).

68. Lee, J.H. et al. ATM directs DNA damage responses and proteostasis via genetically separable pathways. Sci Signal 11 (2018).

69. McVey, M. & Lee, S.E. MMEJ repair of double-strand breaks (director’s cut): deleted sequences and alternative endings. Trends Genet 24, 529–538 (2008).

70. Milella, M. et al. PTEN: Multiple Functions in Human Malignant Tumors. Front Oncol 5, 24 (2015).

71. Ma, J. et al. Inhibition of Nuclear PTEN Tyrosine Phosphorylation Enhances Glioma Radiation Sensitivity through Attenuated DNA Repair. Cancer Cell 35, 504–518 e507 (2019).

