## Supplementary Figures 1-7 for "PTEN mutant NSCLC require ATM to suppress pro-apoptotic signalling and evade radiotherapy"

*Figure S1: PTEN alterations; impact on pathways and radiation resistance*

A. Mean patient survival of various cohorts stratified by PTEN expression. ADC = Adenocarcinoma, SCC = Squamous cell carcinoma, TMB = tumor mutational burden, NSCLC = Non-small-cell lung carcinoma. Generated with the free online tool https://kmplot.com.

B. Schematic representation of CRISPR mediated genome targeting of PTEN exon 1 and 4. Production of lentiviral participles in HEK-293T, infection of BEAS-2B, followed by Blasticidin-selection of positive infected cells 48 hours post infection. Detailed description in Material and Methods.

C. Schematic representation of colony formation assay to assess radiation tolerance in the various cell lines tested. Exposure of cells in suspension to various concentration of ionizing radiation, followed by plating of surviving cells and clonal expansion for 1-2 weeks. Total number of colonies is assessed by manual counting of crystal violet retaining colonies.

D. Survival fraction of parental BEAS-2B cells (black), BEAS-2B $PTEN^{homo}$ (blue) and BEAS-2B $PTEN^{hetero}$ (red) after irradiation with 5 Gy, shown in Figure 1D. Bar graph depicts n=3 experiments and standard deviation.

E. Immunoblot against PTEN and the MAPK pathway proteins BRAF, MEK and phospho-MEK1/2 in parental BEAS-2B, $PTEN^{homo(cloneIII3)}$, or either transduced with a retroviral vector to express GFP (control) or $BRAF^{V600E}$. WT= BEAS $2B^{wt}$ cells; $PTEN^{-/-}$: clone III3 described in 1D. $BRAF^{V600E}$: Epithelial transformed BEAS 2B stable expressing $BRAF^{V600E}$. $BRAF^{V600E}PTEN^{-/-}$: $PTEN^{-/-}$ BEAS 2B cells stable expressing $BRAF^{V600E}$. Actin served as loading control. Representative immunoblots of n=3 experiments.

F. Colony formation assay of BEAS-2B WT (black), $PTEN^{-/-}$ (blue), $BRAF^{v600E}$ (red), $BRAF^{V600E}PTEN^{-/-}$ (pink) after exposure to the indicated doses of ionizing radiation (0-8Gy). SF 2: Surviving fraction at 2 Gy. $D_{25}$: Dose in Gy with 25% survival. Graph represents average values of n=3 experiments and standard deviation.

*Figure S2: Generating and characterizing murine PTEN deficient tumor cell lines*

A. Schematic diagram of the AAV approach of intratracheal infection of mice to induce CRISPR mediated lung tumor formation. KP = simultaneous deletion of *Trp53* and mutation of *KRas* to $KRas^{G12D}$; KPP = simultaneous deletion of *Trp53* and *Pten*, and mutation of *KRas* to $KRas^{G12D}$. 12 weeks post intratracheal infection mice are culled, lungs dissected and primary tumors macroscopically excised and subsequently cultured in standard culture conditions. Please see Material and Methods for further details.

B. Immunoblot against endogenous members of the RTK-MAPK pathway, EGFR, (phospho-)MEK1/2 and (phospho-)ERK1/2, in KP6 and KPP4 cell lines under control conditions or 1h post irradiation with 5 Gy. Quantification of relative protein abundance, normalize to actin as loading control. n/actin: normalized to actin. n/MEK: normalized to MEK. n/ERK normalized to ERK.

C. Immunoblot against the endogenous members of the PI3K/mTOR pathway, PTEN, PIK3CA, (phospho-)AKT, (phospho-)mTOR, (phospho-)S6K and (phospho-)S6, in KP6 and KPP4 cell lines under control conditions or 1h post irradiation with 5 Gy. Quantification of relative protein abundance (actin as loading control). n/actin: normalized to actin. n/AKT: normalized to AKT. n/mTOR: normalized to mTOR. n/S6K: normalized to S6K. n/S6: normalized to S6.

D. Immunoblot of murine KPP4 cells reconstituted with human PTEN via lentiviral transduction, followed by Puromycin selection. KP6 served as PTEN expressing control. Ponceau S (transparent background) to estimate equal loading and proper transfer of protein to the membrane.

*Figure S3: Loss of Pten alters DNA damage signalling pathways in murine NSCLC*

A. Gene set enrichment analyses (GSEA) of KRAS_DN, AKT_MTOR_DN, MYC target genes V1 and G2M Checkpoint gene expression in *$KRas^{G12D}$:Trp53* (KP) relative to *$KRas^{G12D}$:Trp53:Pten* (KPP). (N)ES, normalized enrichment score and p Values are depicted in the table. n=3 each.

B. Gene set enrichment analyses (GSEA) of PACKAGING_OF_TELOMERES, TELOMERE_MAINTENANCE, E2F_TARGETS and REACTIVE_OXYGEN_SPECIES differential gene expression in *$KRas^{G12D}$:Trp53* (KP) relative to *$KRas^{G12D}$:Trp53:Pten* (KPP). (N)ES, normalized enrichment score and p Values are depicted in the table. n=3 each.

C. Representative immunoblot of ionizing radiation (8Gy) pulse chase experiment of endogenous phosphorylated ATR (Thr 1989) and ɣH2ax of KP6 and KPP4 cells post indicated time points. Vinculin as loading control.

*Figure S4: Impact of PI3K/mTOR inhibition in PTEN deficient cells*

A. Schematic representation of the RTK-PI3K-MAPK pathways and the putative therapeutic target structures of PI-103 within these pathways.

B. Schematic overview of the colony forming assay to elucidate the therapeutic benefit of PI-103 treatment. Cells were pre-treated for 3 h with 2 µM PI-103, followed exposure to ionizing irradiation at indicated doses (please see Figure 4), followed by re-seeding 24 h after radiation and counting of crystal violet retaining colonies 1-2 weeks post treatment.

C. Colony formation assay of BEAS-2B WT (black) and BEAS-2B expressing the oncogene BRAF V600E (red), exposed to either DMSO (continuous lines) or 2 µM PI-103 (dashed lines), following treatment with indicated doses of ionizing irradiation. Upon treatment, cells were re-seeded and colonies counted as outlined in S4B. SF2: Surviving fraction at 2 Gy. $D_{25}$: Dose in Gy with 25% survival Graph represents average values of n=3 experiments and standard deviation.

D. Colony formation assay of BEAS-2B WT (black), BEAS-2B *$PTEN^{homozygous}$* (blue) and compound BEAS-2B *$PTEN^{homozygous}$-$BRAF^{V600E}$* compound mutant cells (pink), exposed to either DMSO (continuous lines) or 2 µM PI-103 (dashed lines), following treatment with indicated doses of ionizing irradiation. Upon treatment, cells were re-seeded and colonies counted as outlined in S4B. SF2: Surviving fraction at 2 Gy. $D_{25}$: Dose in Gy with 25% survival. Graph represents average values of n=3 experiments and standard deviation.

*Figure S5: Impact of ATM inhibition in PTEN deficient cells*

A. Dose response and colony formation ability of murine *Pten* proficient KP6 (black) and *Pten* deficient KPP4 (blue) cells following treatment with the ATM inhibitor AZD 1390 at indicated concentrations for 27 h. Graph represents average values of n=3 experiments and standard deviation.

B. Dose response and colony formation ability of PTEN deficient (blue) and $PTEN^{homozygous}$-$BRAF^{V600E}$ compound mutant BEAS-2B cells (pink) following treatment with the ATM inhibitor KU-60019 at indicated concentrations for 27 h. Graph represents average values of n=3 experiments and standard deviation.

C: Immunoblot against endogenous ATM, phospho-S1987-ATM, AKT and phospho-S473-AKT of KP6 and KPP4 cells pre-treated for 3 h with either DMSO or indicated concentrations of the ATM inhibitor AZD 1390, following IR with 8 Gy and collection of samples 30 min post IR. DMSO served as solvent control. Actin served as loading control. Representative immunoblots of n=3 experiments.

D. Colony formation assay of BEAS-$2B^{wildtype}$ (black) and BEAS-2B $PTEN^{homozygous}$ cells (blue), pre-treated for 3 h with 0.3 µM KU-60019 (dashed lines) or DMSO as control solvent (continuous lines), following exposure to indicated doses of IR. Treated cells were re-seeded and colony forming capacity assessed as outlined in Figure S4B. SF 2: Surviving fraction at 2 Gy. $D_{25}$: Dose in Gy with 25% survival. Graph represents average values of n=3 experiments and standard deviation.

E. Colony formation assay of BEAS-2B $PTEN^{homozygous}$ (blue) and compound BEAS-2B $PTEN^{homozygous}$-$BRAF^{V600E}$ compound mutant cells (pink), pre-treated for 3 hours with 3 µM KU-60019 (dashed lines) or DMSO as control solvent (continuous lines), following exposure to indicated doses of IR. Treated cells were re-seeded and colony forming capacity assessed as outlined in Figure S4B. SF 2: Surviving fraction at 2 Gy. $D_{25}$: Dose in Gy with 25% survival. Graph represents average values of n=3 experiments and standard deviation.

F. Colony formation assay of murine *Pten* proficient KP6 (black) and *Pten* deficient KPP4 (blue) cells pre-treated for 3 h with 3 µM AZD 1390 (dashed lines) or DMSO as control (continuous lines), following exposure to indicated doses of IR. Treated cells were re-seeded and colony forming capacity assessed as outlined in Figure S4B. SF 2: Surviving fraction at 2 Gy. $D_{25}$: Dose in Gy with 25% survival. Graph represents average values of n=3 experiments and standard deviation.

*Figure S6: Multilevel proteomics show differential apoptosis signaling*

A. Enrichment map of Reactome pathways differentially phosphorylated between KP and KPP cells under unperturbed conditions (Log2 FC >0.5, P value < 0.05). Related pathways are connected by edges. Node coloring corresponds to ReactomeFI functional enrichment score. All pathways shown are significantly enriched with an FDR < 0.05.

B. Lollipop plot showing pathway enrichment results of genes differentially regulated in phosphorylation upon combinatorial treatment between KP6 and KPP4 cell lines (log2 FC difference >0.5). Size and coloring according to the number of genes found differentially regulated. Additionally, the enrichment FDR is given on the y axis. Pathways have been sorted according to FDR

C. APC-Annexin V/DAPI staining of *Pten* proficient KP6 and *Pten* deficient KPP4 cells pre-treated for 3 h with 3 µM KU-60019, following irradiation with 8 Gy. Cell viability was assessed 24h, 48h, 72h and 96h post irradiation.

D. Cell survival of KP6 and KPP4 cells 48 hours post exposure to 8 Gy and 3 µM KU-60019 or exposure to 5 µM Camptothecin (CPT). CPT served as positive control for the cell survival FACS analysis, as shown in Figure 5. Prior to cell viability marker staining, cells were trypsinised and pooled with the whole cultivating medium to collect dead/dying cells. The graph shows the apoptotic fraction (Annexin $V^{positive}$/$DAPI^{positive}$) of KP6 and KPP4 cells collected at indicated timepoints.

*Figure S7: Tumor slice culture response to KU-60019 treatment and radiation*

A. Schematic representation of the organotypic lung slice culture. GFP+ murine lung cancer cell lines KP6 and KPP4 were orthotopically re-transplanted. Upon engraftment, lungs were excised and whole lung lobes sectioned using a Leica V1200S vibratome. Cultured lung slices were cultured in the presence of either DMS0 or 3 µM ATM inhibitor KU-60019, following ionizing irradiation with 8 Gy. Treated tissue slices were longitudinally cultured and pictures of same tumor sites were taken for a total of 8 days.

B. Representative images of GFP-positive KP6 and KPP4 tumors upon treatment with DMSO, 3 µM ATM inhibitor KU-60019, IR of 8 Gy or combinatorial treatment. Relative GFP signal intensity was measured by brightfield microscopy over a total of 7-8 days. Inlay images show representative individual tumor bearing lung slice cultures at days 0/1, 2, 4, 6 and at endpoint day 7/8. Tumors are indicated by dashed yellow line.

A

median patient survival

| *Analysis* | *PTEN^low^* | *PTEN^high^* |
|---|---|---|
| ADC | 56 | 175 |
| SCC | 42 | 64 |
| Female | 86 | 120 |
| Male | 48 | 65 |
| TMB low | 46 | 58 |
| TMB high | 39 | 53 |
| Neversmoker | 76 | n.a. |
| Smoker | 69 | 96 |
| NSCLC | 57 | 81 |
| Radiotherapy | 12 | 27 |

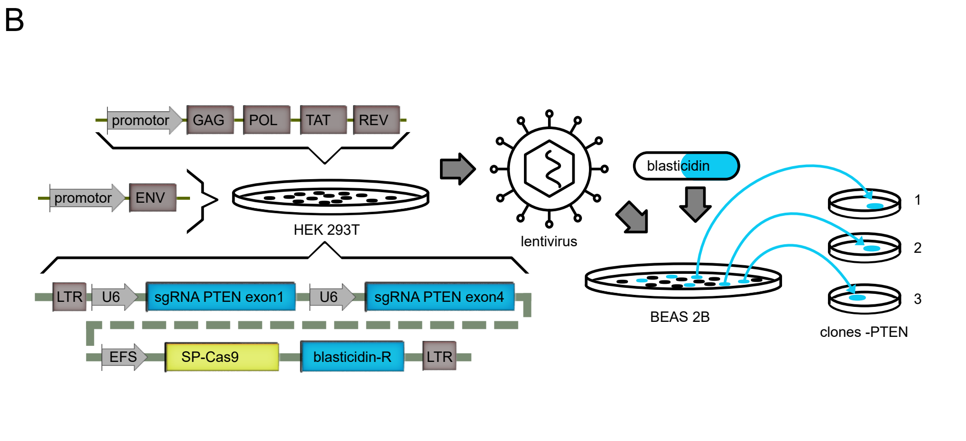


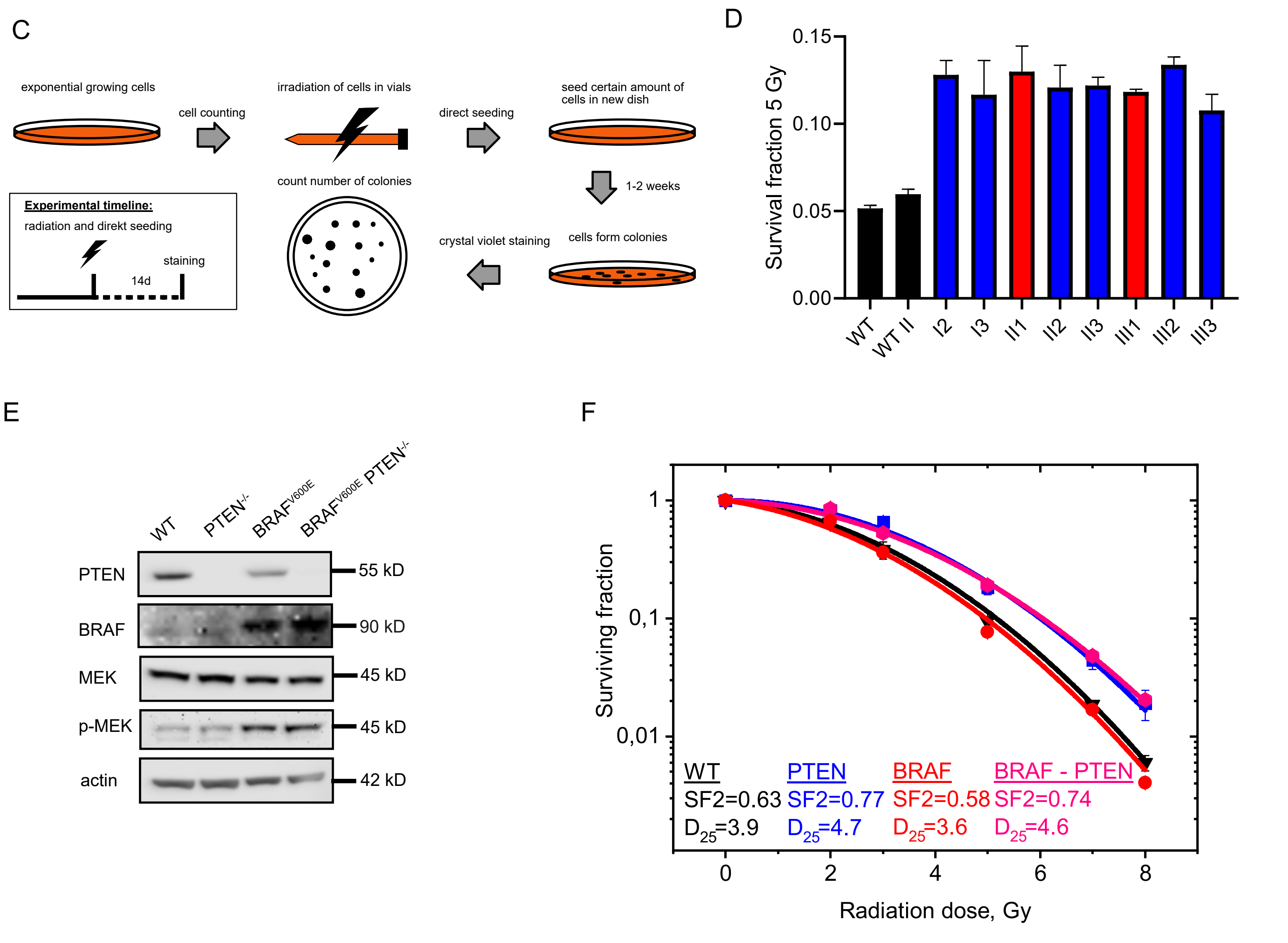

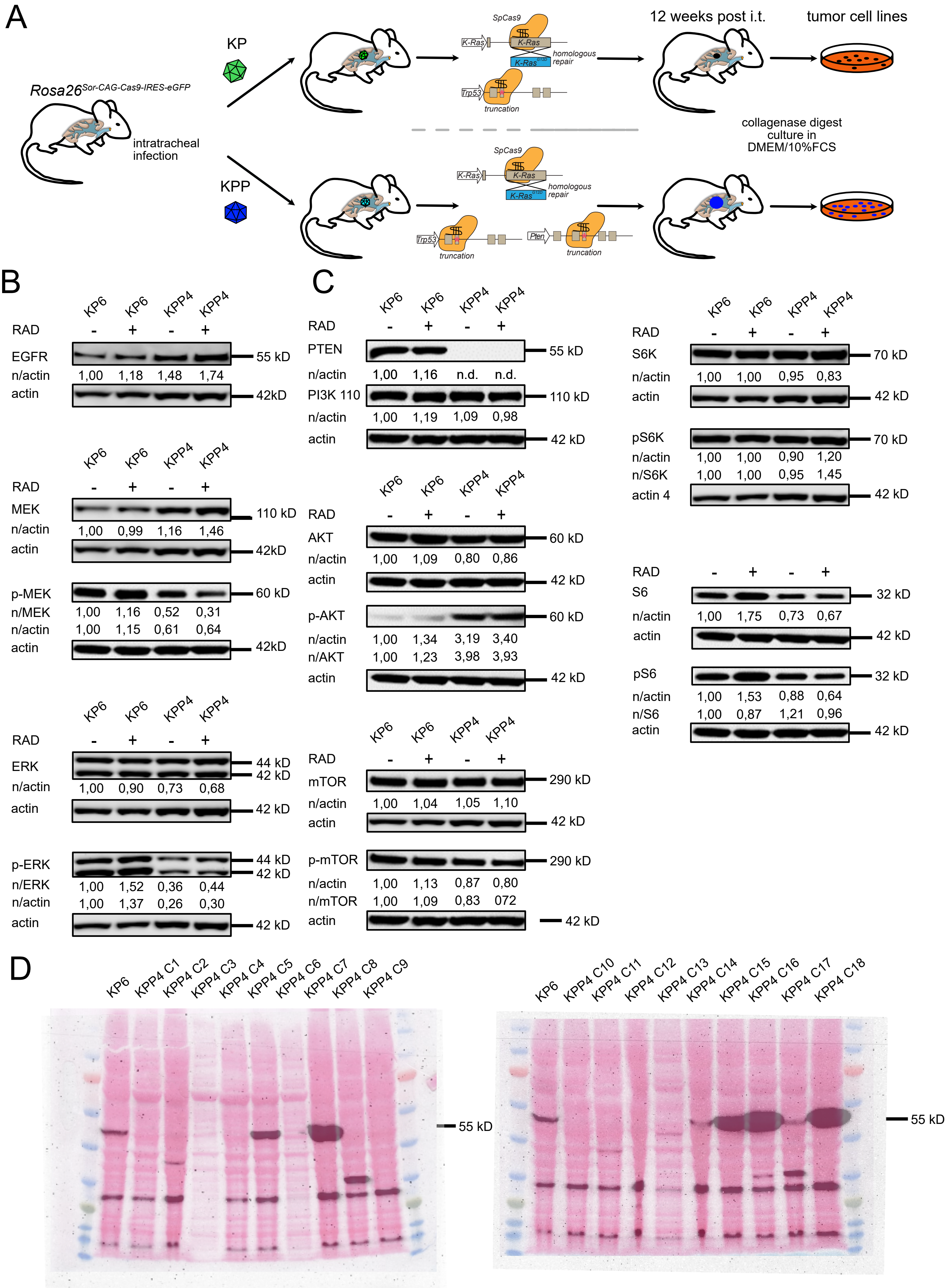
A
KP
KPP
Rosa26Sor-CAG-Cas9-IRES-eGFP
intratracheal infection
SpCas9
K-Ras
K-Ras G12D
homologous repair
Trp53
Pten
truncation
12 weeks post i.t.
tumor cell lines
collagenase digest
culture in
DMEM/10%FCS
B
KP6 KP6 KPP4 KPP4
RAD - + - +
EGFR 55 kD
n/actin 1,00 1,18 1,48 1,74
actin 42kD
MEK 110 kD
n/actin 1,00 0,99 1,16 1,46
actin 42kD
p-MEK 60 kD
n/MEK 1,00 1,16 0,52 0,31
n/actin 1,00 1,15 0,61 0,64
actin 42kD
ERK 44 kD 42 kD
n/actin 1,00 0,90 0,73 0,68
actin 42 kD
p-ERK 44 kD 42 kD
n/ERK 1,00 1,52 0,36 0,44
n/actin 1,00 1,37 0,26 0,30
actin 42 kD
C
PTEN 55 kD
n/actin 1,00 1,16 n.d. n.d.
PI3K 110 110 kD
n/actin 1,00 1,19 1,09 0,98
actin 42 kD
AKT 60 kD
n/actin 1,00 1,09 0,80 0,86
actin 42 kD
p-AKT 60 kD
n/actin 1,00 1,34 3,19 3,40
n/AKT 1,00 1,23 3,98 3,93
actin 42 kD
mTOR 290 kD
n/actin 1,00 1,04 1,05 1,10
actin 42 kD
p-mTOR 290 kD
n/actin 1,00 1,13 0,87 0,80
n/mTOR 1,00 1,09 0,83 072
actin 42 kD
S6K 70 kD
n/actin 1,00 1,00 0,95 0,83
actin 42 kD
pS6K 70 kD
n/actin 1,00 1,00 0,90 1,20
n/S6K 1,00 1,00 0,95 1,45
actin 4 42 kD
S6 32 kD
n/actin 1,00 1,75 0,73 0,67
actin 42 kD
pS6 32 kD
n/actin 1,00 1,53 0,88 0,64
n/S6 1,00 0,87 1,21 0,96
actin 42 kD
D
KP6 KPP4 C1 KPP4 C2 KPP4 C3 KPP4 C4 KPP4 C5 KPP4 C6 KPP4 C7 KPP4 C8 KPP4 C9
55 kD
KP6 KPP4 C10 KPP4 C11 KPP4 C12 KPP4 C13 KPP4 C14 KPP4 C15 KPP4 C16 KPP4 C17 KPP4 C18
55 kD

A

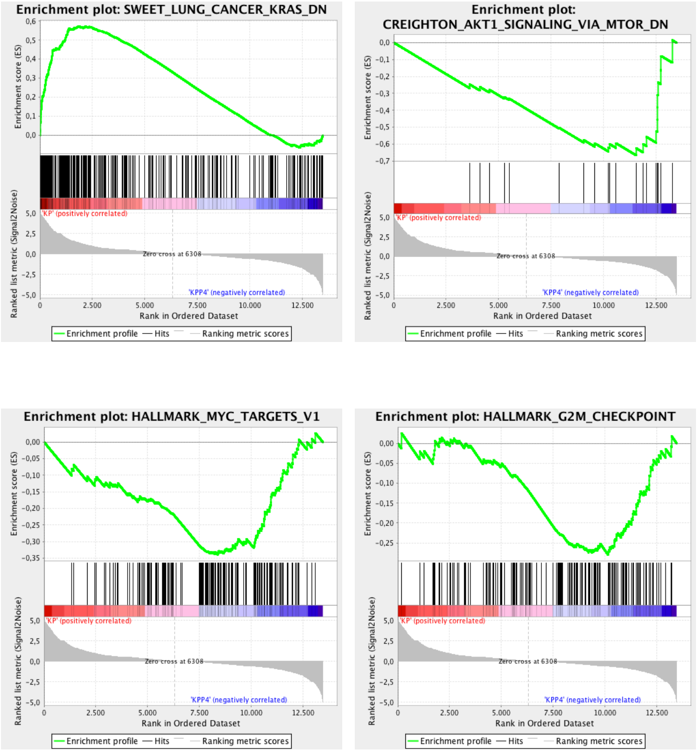


| *Geneset* | *(N)ES* | *pValue* |
|---|---|---|
| KRAS_DN | 2.15 | <0.0005 |
| AKT_MTOR_DN | -1.91 | <0.0005 |
| MYC_TARGET_V1 | -1.42 | <0.0005 |
| G2M | -1.16 | 0.07 |

B

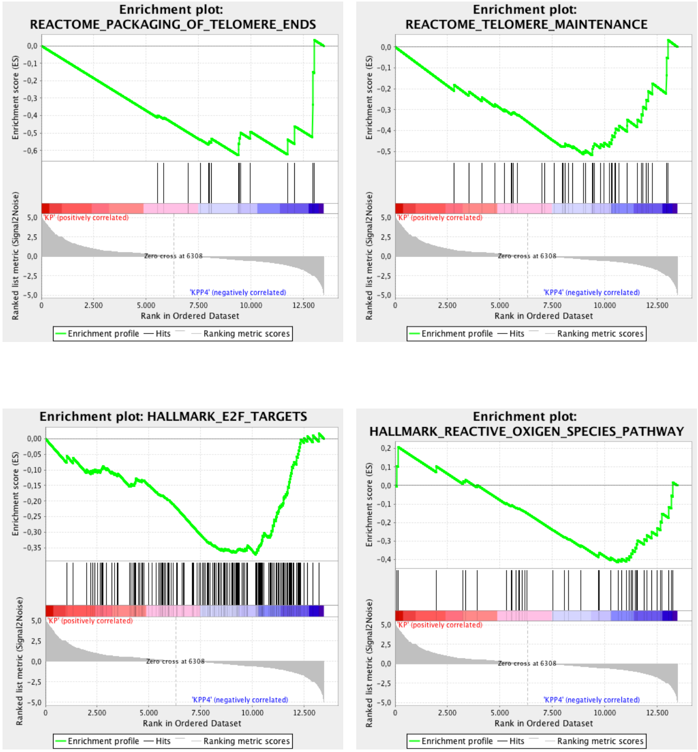


| *Geneset* | *(N)ES* | *pValue* |
|---|---|---|
| PACK._TELOMERE | 1.74 | <0.0005 |
| TELOMERE_MAINT. | -2.45 | <0.0005 |
| E2F | -1.42 | <0.0005 |
| ROS_PATHWAY | -1.35 | 0.05 |

C

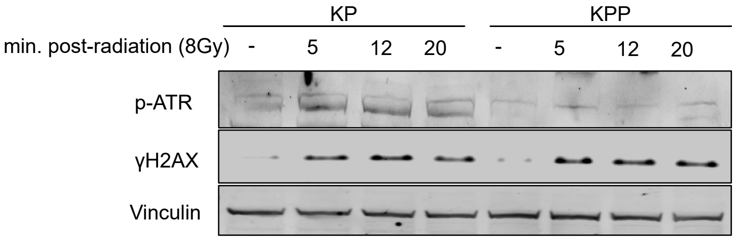


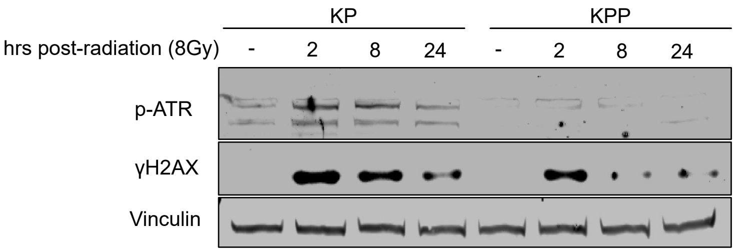

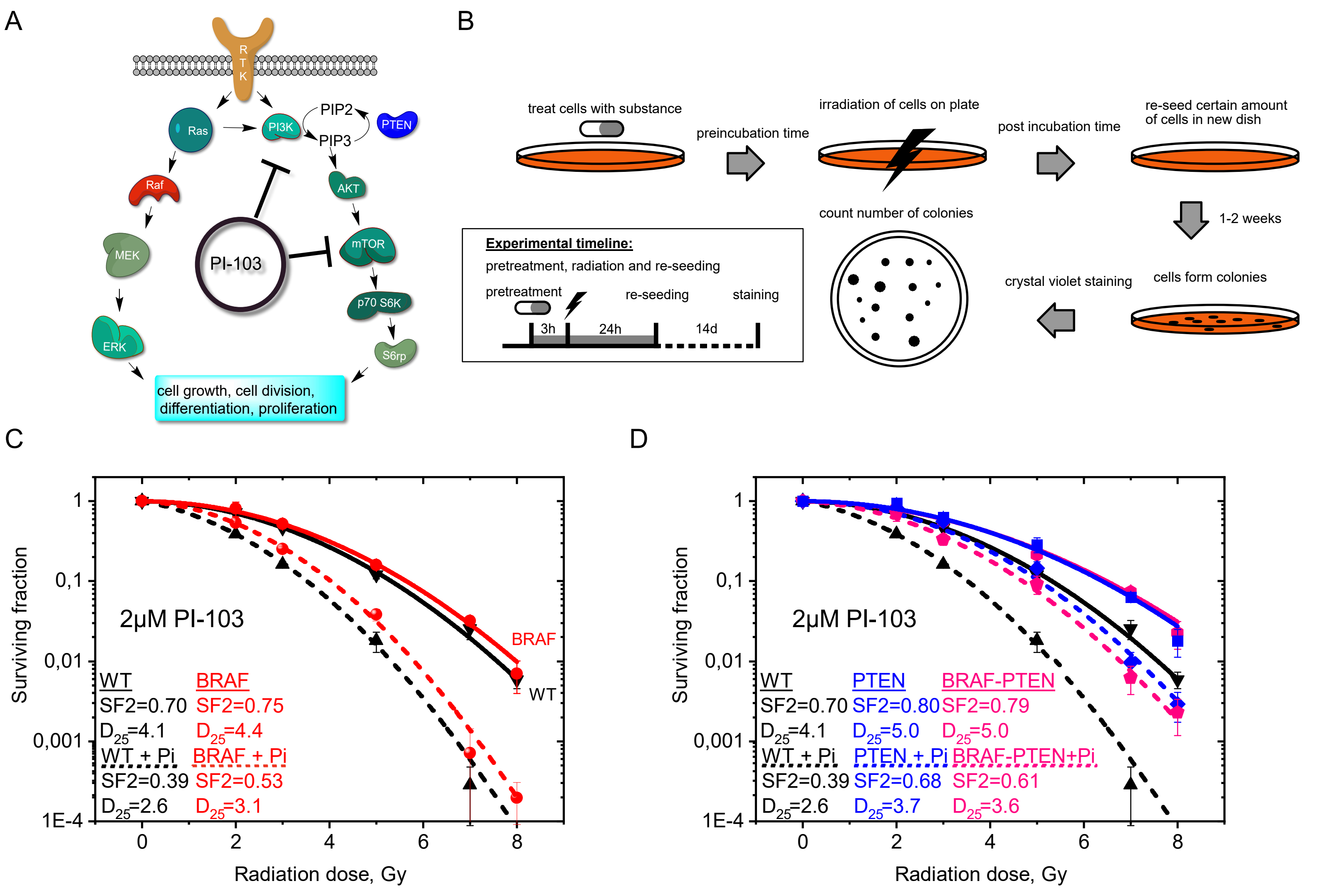

A
RTK
Ras
PI3K
PIP2
PIP3
PTEN
Raf
AKT
MEK
PI-103
mTOR
p70 S6K
ERK
S6rp
cell growth, cell division, differentiation, proliferation
B
treat cells with substance
preincubation time
irradiation of cells on plate
post incubation time
re-seed certain amount of cells in new dish
1-2 weeks
count number of colonies
Experimental timeline:
pretreatment, radiation and re-seeding
pretreatment
re-seeding
staining
3h
24h
14d
crystal violet staining
cells form colonies
C
2µM PI-103
Surviving fraction
Radiation dose, Gy
BRAF
WT
WT
SF2=0.70
D25=4.1
BRAF
SF2=0.75
D25=4.4
WT + Pi
SF2=0.39
D25=2.6
BRAF + Pi
SF2=0.53
D25=3.1
D
2µM PI-103
Surviving fraction
Radiation dose, Gy
WT
SF2=0.70
D25=4.1
PTEN
SF2=0.80
D25=5.0
BRAF-PTEN
SF2=0.79
D25=5.0
WT + Pi
SF2=0.39
D25=2.6
PTEN + Pi
SF2=0.68
D25=3.7
BRAF-PTEN+Pi
SF2=0.61
D25=3.6

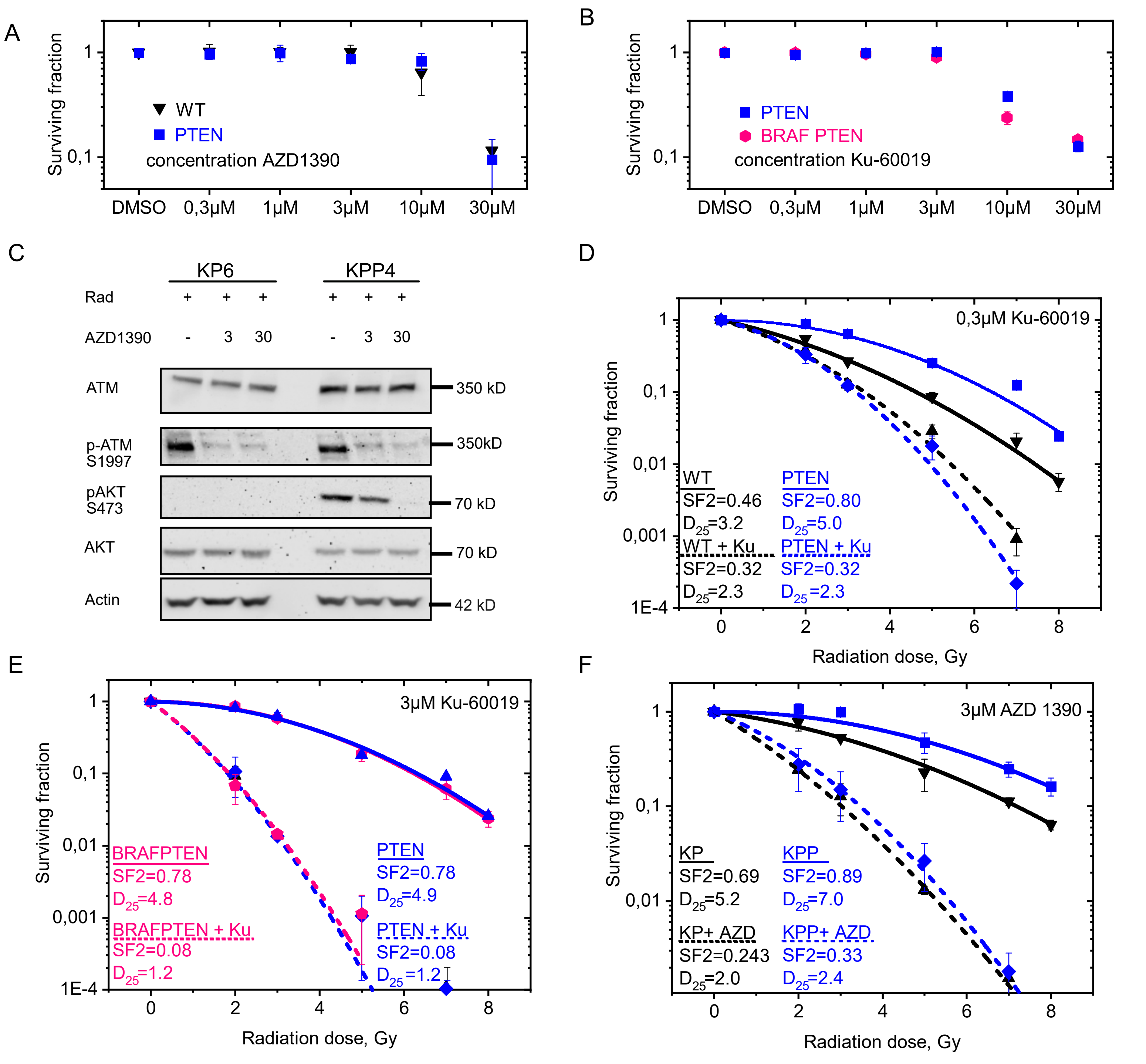
A
Surviving fraction
1
0,1
WT
PTEN
concentration AZD1390
DMSO
0,3µM
1µM
3µM
10µM
30µM
B
Surviving fraction
PTEN
BRAF PTEN
concentration Ku-60019
C
KP6
KPP4
Rad
AZD1390
ATM
p-ATM S1997
pAKT S473
AKT
Actin
350 kD
350kD
70 kD
70 kD
42 kD
D
0,3µM Ku-60019
WT
SF2=0.46
D25=3.2
PTEN
SF2=0.80
D25=5.0
WT + Ku
SF2=0.32
D25=2.3
PTEN + Ku
SF2=0.32
D25=2.3
Radiation dose, Gy
E
3µM Ku-60019
BRAFPTEN
SF2=0.78
D25=4.8
PTEN
SF2=0.78
D25=4.9
BRAFPTEN + Ku
SF2=0.08
D25=1.2
PTEN + Ku
SF2=0.08
D25=1.2
Radiation dose, Gy
F
3µM AZD 1390
KP
SF2=0.69
D25=5.2
KPP
SF2=0.89
D25=7.0
KP+ AZD
SF2=0.243
D25=2.0
KPP+ AZD
SF2=0.33
D25=2.4
Radiation dose, Gy

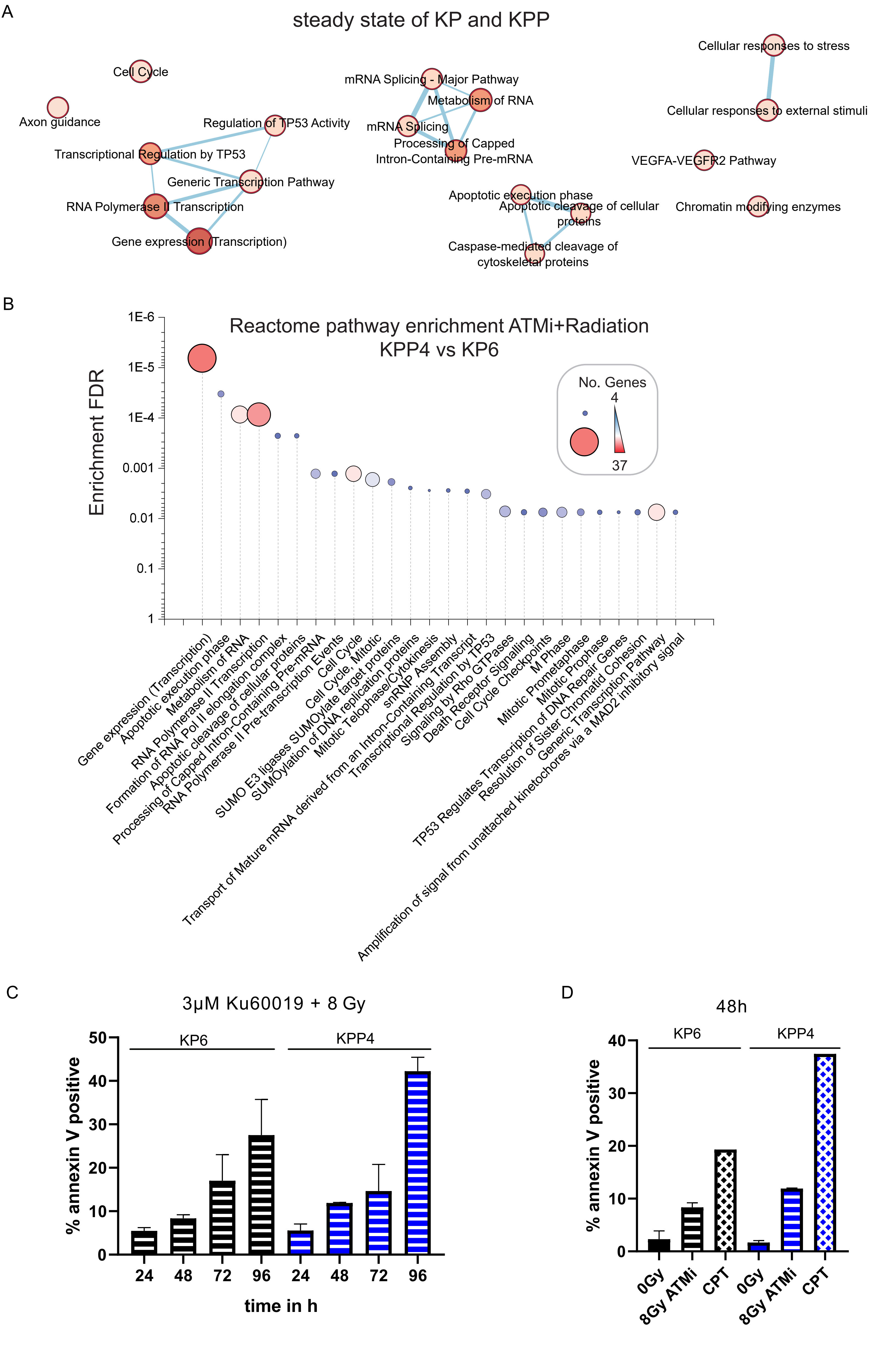
A
steady state of KP and KPP
Cell Cycle
Axon guidance
Regulation of TP53 Activity
Transcriptional Regulation by TP53
Generic Transcription Pathway
RNA Polymerase II Transcription
Gene expression (Transcription)
mRNA Splicing - Major Pathway
Metabolism of RNA
mRNA Splicing
Processing of Capped Intron-Containing Pre-mRNA
Apoptotic execution phase
Apoptotic cleavage of cellular proteins
Caspase-mediated cleavage of cytoskeletal proteins
Cellular responses to stress
Cellular responses to external stimuli
VEGFA-VEGFR2 Pathway
Chromatin modifying enzymes
B
Reactome pathway enrichment ATMi+Radiation
KPP4 vs KP6
Enrichment FDR
1E-6
1E-5
1E-4
0.001
0.01
0.1
1
No. Genes
4
37
Gene expression (Transcription)
Apoptotic execution phase
Metabolism of RNA
RNA Polymerase II Transcription
Formation of RNA Pol II elongation complex
Apoptotic cleavage of cellular proteins
Processing of Capped Intron-Containing Pre-mRNA
RNA Polymerase II Pre-transcription Events
Cell Cycle
Cell Cycle, Mitotic
SUMO E3 ligases SUMOylate target proteins
SUMOylation of DNA replication proteins
Mitotic Telophase/Cytokinesis
snRNP Assembly
Transport of Mature mRNA derived from an Intron-Containing Transcript
Transcriptional Regulation by TP53
Signaling by Rho GTPases
Death Receptor Signalling
Cell Cycle Checkpoints
M Phase
Mitotic Prometaphase
Mitotic Prophase
TP53 Regulates Transcription of DNA Repair Genes
Resolution of Sister Chromatid Cohesion
Generic Transcription Pathway
Amplification of signal from unattached kinetochores via a MAD2 inhibitory signal
C
3µM Ku60019 + 8 Gy
KP6
KPP4
% annexin V positive
0
10
20
30
40
50
24
48
72
96
24
48
72
96
time in h
D
48h
KP6
KPP4
% annexin V positive
0
10
20
30
40
0Gy
8Gy ATMi
CPT
0Gy
8Gy ATMi
CPT

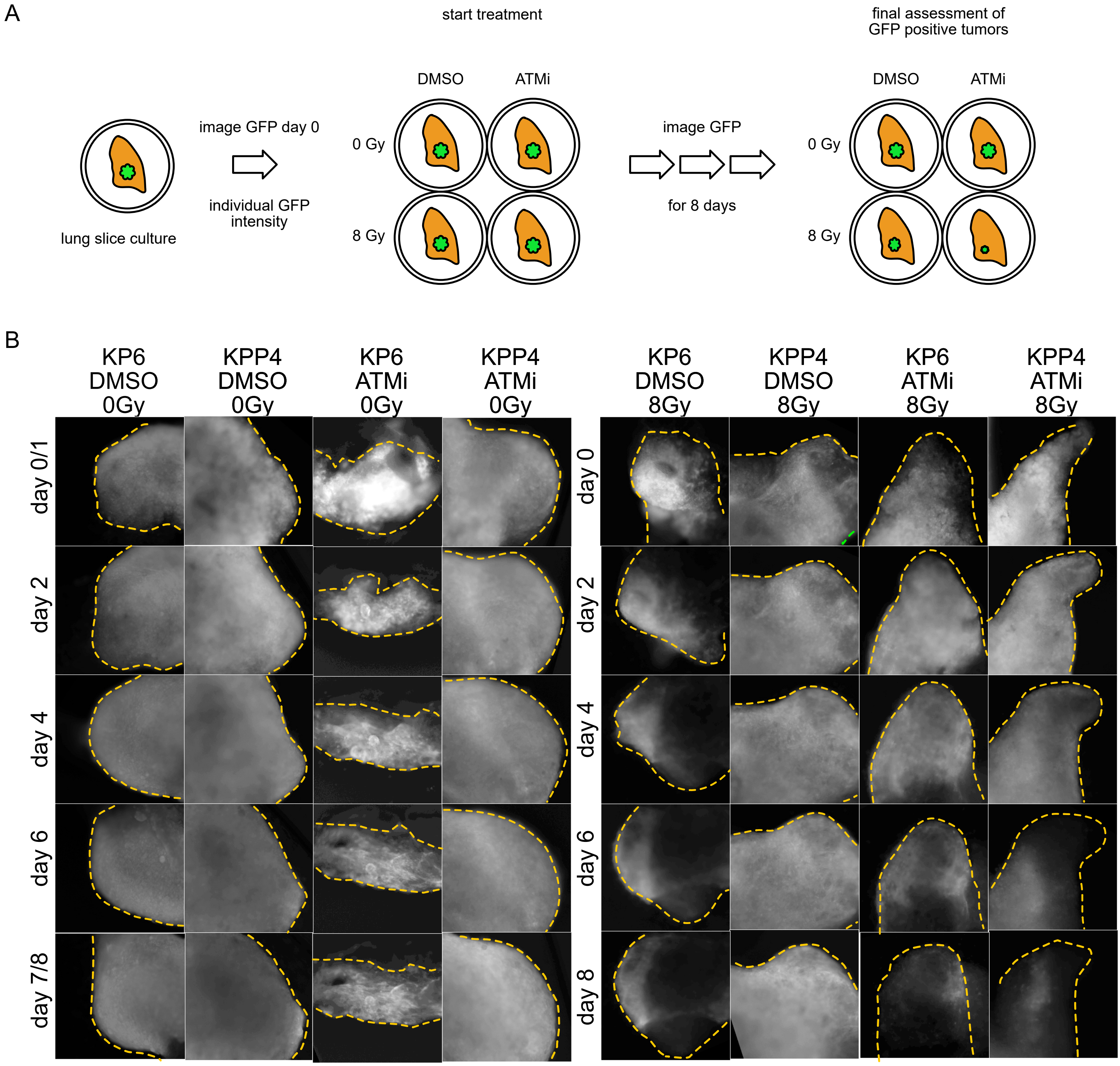
A
start treatment
final assessment of
GFP positive tumors
lung slice culture
image GFP day 0
individual GFP
intensity
DMSO
ATMi
0 Gy
8 Gy
image GFP
for 8 days
B
KP6 DMSO 0Gy
KPP4 DMSO 0Gy
KP6 ATMi 0Gy
KPP4 ATMi 0Gy
KP6 DMSO 8Gy
KPP4 DMSO 8Gy
KP6 ATMi 8Gy
KPP4 ATMi 8Gy
day 0/1
day 0
day 2
day 4
day 6
day 7/8
day 8
