## Supplementary material for "PTEN mutant NSCLC require ATM to suppress pro-apoptotic signalling and evade radiotherapy": STAR Methods

| **1^st^ Antibodies** | **Company** | **Identifier** | **RRID** |
| --- | --- | --- | --- |
| Monoclonal rabbit anti-pAKT Ser473 | Cell signalling | 4060 | AB_2315049 |
| Polyclonal rabbit anti-AKT | Cell signalling | 9272 | AB_329827 |
| Polyclonal rabbit anti-p-ATM (ser1981) | R&D Sytems | AF1655 | AB_2062935 |
| Monoclonal rabbit anti-ATM | Cell signalling | 2873 | AB_2062659 |
| Monoclonal rabbit anti p Histone-H2A.X(ser139) | Cell signalling | 9718 | AB_2118009 |
| Polyclonal rabbit anti p-ATR (ser428) | Cell signalling | 2853 | AB_2290281 |
| Monoclonal rabbit anti-EGFR | Bimake | A5858 |  |
| Monoclonal mouse anti-S6 | Cell signalling | 2317 | AB_2238583 |
| Polyclonal rabbit anti-p-S6 | Cell signalling | 2215 | AB_331682 |
| Polyclonal rabbit anti-PTEN | Cell signalling | 9552 | AB_10694066 |
| Monoclonal rabbit anti-S6K | Bimake | A5512 |  |
| Monoclonal rabbit anti-pS6K | Bimake | A5033 |  |
| Polyclonal rabbit anti-ERK | Cell signalling | 9102 | AB_330744 |
| Monoclonal rabbit anti-pERK | Cell signalling | 4370 | AB_2315112 |
| Monoclonal rabbit anti-MEK | Cell signalling | 9126 | AB_331778 |
| Monoclonal rabbit anti-pMEK | Cell signalling | 9154 | AB_2138017 |
| Monoclonal mouse anti-Actin | Calbiochem | CP01 | AB_566293 |
| Monoclonal rabbit anti-BRAF | Bimake | A5104 |  |

| **­2^nd^ Antibodies** | **Company** | **Identifier** | **RRID** |
| --- | --- | --- | --- |
| Polyclonal goat anti-mouse HRP | DAKO | P0447 | AB_2617137 |
| Polyclonal goat anti rabbit HRP | DAKO | P0048 | AB_2617138 |
| SuperBoost™ Goat anti-Mouse Poly HRP | ThermoFisher | B40961 |  |
| SuperBoost™ Goat anti-Rabbit Poly HRP | ThermoFisher | B40962 |  |

| **Bacterial Strains** | **Company** | **Identifier** |
| --- | --- | --- |
| **DH5α** F- endA1 glnV44 thi-1 recA1 relA1 gyrA96 deoR nupG Φ80dlacZΔM15 Δ(lacZYA-argF)U169, hsdR17(rK-mK+), λ– | ThermoFisher | 18263012 |
| **Chemicals and Commercial Assays** | **Company** | **Identifier** |
| Sigma™  Dulbecco’s Modified Eagle Medium (DMEM), high glucose | Sigma-Aldrich | D6429 |
| Sigma™ Trypsin-EDTA (0.5%), Phenol Red | Sigma-Aldrich | T4049 |
| Fetal Bovine Serum (FCS) | Sigma-Aldrich | 12103C |
| Penicillin-Streptomycin | Sigma-Aldrich | P4333 |
| Dulbecco’s Phosphate Buffered Saline | Sigma-Aldrich | D8537 |
| Polybrene | Sigma-Aldrich | TR-1003 |
| Polyethylenimine, Linear, MW 25000, Transfection Grade (PEI 25K) | Polysciences | 23966-1 |
| Dimethyl sulfoxide (DMSO) | Sigma-Aldrich | D8418 |
| Ethanol (Etoh) | Carl Roth | 5054.6 |
| Nuclease-free water | Merck | 3098 |
| KU-60019 | Selleckchem | S1570 |
| AZD 1390 | Selleckchem | S8680 |
| PI-103 | Selleckchem | S1038 |
| Blasticidin | Invivogen | ant-bl-1 |
| Puromycin | Invivogen | ant-pr-1 |
| Propidium Iodide (PI) | Sigma-Aldrich | 11348639001 |
| Protease Inhibitor Cocktail | Roche | 4693159001 |
| Bovine serum albumine (BSA) | Merck | 810683 |
| RIPA Lysis Buffer | Homemade |  |
| 4′,6-diamidino-2-phenylindole (DAPI) | ThermoFisher | D1306 |
| Phosphatase-inhibitor-cocktail | Bimake | B15001 |
| Phosphate-buffered saline (PBS) | Homemade |  |
| 2-Propanol/ Isopropanol | ROTH | AE73.2 |
| Adenosintriphosphat (ATP) | Jena Bioscience | NU-1010-10G |
| Agarose | ROTH | 3810.4 |
| Ampicillin (Amp) | ROTH | HP62.2 |
| Bovine serum albumine (BSA) | Merck Millipore | 810683 |
| Mowiol® 40-88 | Sigma-Aldrich | 324590 |
| Dithiothreitol (DTT) | Sigma-Aldrich | D9779 |
| Eosin | Sigma | E4009 |
| Hematoxylin | Sigma | H3136 |
| PonceauS solution | Sigma | P7170 |
| Crystal violet | Sigma | C3886 |
| Natrium chloride (NaCl) | AppliChem | A2942,1000 |
| Neutrally buffered formalin (NBF) | Thermo Fisher | 5700TS |
| Tris-HCl | ROTH | 9090.5 |
| TritonX100 | ROTH | 3051.3 |
| Xylene | Sigma | 534056 |
| H_2_O_2_ 30% | ROTH | 8070.2 |
| Methanol (MeOH) | ROTH | 0082.3 |
| Acetic Acid | Chemsolute | 2289.2500 |
| peq GOLD Trifast | VWR (Peqlab brand) | 30-2010 |
| Deoxynucleotidetriphosphates (dNTPs) Mix | Promega | U151A |
| Random Hexamer Primer | ThermoFisher | SO142 |
| RiboLock RNase Inhibitor | ThermoFisher | EO0381 |
| ReliaPrep™ RNA Cell Miniprep System Protocol | Promega | TM370 |
| NEBNext® Ultra™RNA Library Prep Kit for Illumina | New England Biolabs (NEB) | NEB #E7530S |
| NEBNext® Multiplex Oligos for Illumina® (Dual Index Primers Set 1) | New England Biolabs (NEB) | NEB #E7600S |
| NEBNext® Poly(A) mRNA Magnetic Isolation Module | New England Biolabs (NEB) | NEB #E7490S |
| APC Apoptosis Detektion Kit | Biolegend | 640932 |
| Annexin V Binding Buffer | Biolegend | 42201 |
| NuPAGE Transfer Buffer | Novex | MP0006-1 |
| Tris-Acetat SDS Running Buffer | Novex | LA0041 |
| MES SDS Running Buffer | Novex | NP0002 |
| MOPS SDS Running Buffer | Novex | NP0001 |
| Nu-PAGE 4-12% Bis-Tris-Gel | Novex | NP0322Box |
| Nu-PAGE 3-8% Tris-Acetat-Gel | Novex | EA03752Box |
| Nitrocellulose Membrane | Novex | LC2000 |
| Nu-PAGE LDS sample Buffer | Novex | NP0007 |
| Nu-PAGE sample reducing Agent | Novex | NP0009 |
| Nu-PAGE antioxidant | Novex | NP0005 |
| **Cell lines**  **Cell** | **Company** | **Identifier** |
| Human: HEK 293T | ATCC | ATCC® CRL-11268™ |
| Human: BEAS-2B | ATCC | ATCC® CRL-9609 |
| Mouse: KP | Primary tumors |  |
| Mouse: KPP | Primary tumors |  |
| **Experimental Models: Organisms/Strains Cell** | **Company** | **Identifier** |
| B6(C)-Gt(ROSA)26Sor^em1.1(CAG-cas9*,-EGFP)Rsky^/J | The Jackson laboratory | Stock No: 028555 |
| C57BL/6J | The Jackson Laboratory | Stock No: 000664 |
| **Oligonucleotides** | **Sequence** | **Company** |
| sgRNA murine Pten 1 for | CACCGTGTGCATATTTATTGCATCG | Sigma |
| sgRNA murine Pten 1 rev | AAACCGATGCAATAAATATGCACAC | Sigma |
| sgRNA human PTEN exon 1 for | CACCGCAGCCGCAGAAATGGATAC | Sigma |
| sgRNA human PTEN exon 1 rev | AAACCCAAATTTAATTGCAGAGGTc | Sigma |
| sgRNA human PTEN exon 4 for | CACCGACCTCTGCAATTAAATTTGG | Sigma |
| sgRNA human PTEN exon 4 rev | AAACCCAAATTTAATTGCAGAGGTC | Sigma |
| sgRNA murine Kras #1 for | CACCGACTGAGTATAAACTTGTGG | Sigma |
| sgRNA murine Kras #1 rev | AAACCCACAAGTTTATACTCAGTC | Sigma |
| sgRNA murine Trp53 #1 for | CACCGATGGTGGTATACTCAGAGC | Sigma |
| sgRNA murine Trp53 #1 rev | AAACGCTCTGAGTATACCACCATC | Sigma |
| KrasG12D repair template for | TTTTGTGTAAGCTTTGGTAACTCCATGTATTTTTATTAAGTGTT | Sigma |
| KrasG12D repair template rev | GAGCTTATCGATACCGTCGACACACCCAGTTTAAAGCCTTGGAA | Sigma |
| **Recombinant DNA** | **Company/Source** | **Identifier** |
| AAV:ITR-U6-sgRNA(Kras)-U6-sgRNA(p53)-U6-sgRNA(Pten)-pEFS-2A-mCherry-shortPA-KrasG12D_HDRdonor-ITR | This publication | N/A |
| pSICO-sgPTENExon1-sgPTENExon4-spCas9-p2A-Blasti | This publication | N/A |
| pSico | pSICO was a gift from Tyler Jacks (Addgene plasmid # 11578 ; http://n2t.net/addgene: 11578 ; RRID:Addgene 11578) | 11578 |
| lentiCas9-Blast | lentiCas9-Blast was a gift from Feng Zhang (Addgene plasmid # 52962 ; http://n2t.net/addgene:52962 ; RRID:Addgene_52962) | 52962 |
| psPAX2 | psPAX2 was a gift from Didier Trono (Addgene plasmid # 12260 ; http://n2t.net/addgene:12260 ; RRID:Addgene_12260) | Addgene plasmid # 12260 |
| pMD2G | pMD2.G was a gift from Didier Trono (Addgene plasmid # 12259 ; http://n2t.net/addgene:12259 ; RRID:Addgene_12259) | Addgene plasmid # 12259 |
| pHelper | Cell Biolabs, INC. | VPK-400-DJ |
| pAAV2/8 | AAV2/8 was a gift from James M. Wilson (Addgene plasmid # 17544 ; http://n2t.net/addgene:112864 ; RRID:Addgene_112864) | Addgene plasmid # 112864 |
| pAAV-DJ Vector | Cell Biolabs, INC. | VPK-420-DJ |
| pBabe B-Raf V600E | pBabe B-Raf V600E was a gift from Channing Der (Addgene plasmid # 17544 ; http://n2t.net/addgene:17544 ; RRID:Addgene_17544) | Addgene plasmid # 17544 |
| gag/pol | gag/pol was a gift from Tannishtha Reya (Addgene plasmid # 14887 ; http://n2t.net/addgene:14887 ; RRID:Addgene_14887) | 14887 |
| pCMV-VSV-G | pCMV-VSV-G was a gift from Bob Weinberg (Addgene plasmid # 8454 ; http://n2t.net/addgene:8454 ; RRID:Addgene_8454) | 8454 |
| **Experimental Models: Organisms/Strains Cell** | **Company** | **Identifier** |
| B6(C)-Gt(ROSA)26Sor^em1.1(CAG-cas9*,-EGFP)Rsky^/J | The Jackson laboratory | Stock No: 028555 |
| C57BL/6J | The Jackson Laboratory | Stock No: 000664 |
| **Software and Algorithm** | **Company/Source** |  |
| cBioportal | https://www.cbioportal.org |  |
| GEPIA and GEPIA2 | http://gepia.cancer-pku.cn |  |
| KM-plotter | http://kmplot.com/analysis/ |  |
| Operetta Imaging | Perkin Elmar |  |
| BoxPlotR | http://shiny.chemgrid.org/boxplotr/ |  |
| Excel | Microsoft |  |
| Affinity Desgner | https://affinity.serif.com/es/designer/ |  |
| Image Studio | Licor |  |
| Panther Classification system | http://pantherdb.org |  |
| AATBIO IC50 calculator | https://www.aatbio.com/tools/ic50-calculator |  |
| GraphPad Software | GraphPad Software, Inc. |  |
| OriginPro | OriginLab Corporation |  |
| ImageJ | National Insistute of Health |  |
| Primerx | http://www.bioinformatics.org/primerx/cgi-bin/DNA_1.cgi |  |
| ROC Plotter | http://www.rocplot.org/ |  |
| Pannoramic Case Viewer | 3dHistech |  |
| R2: Genomics Analysis and Visualization Platform | http://r2.amc.nl |  |
| UCSC Xena | https://ucsc-xena.gitbook.io/project/ |  |
| GenerateFastq v1.1.0.64 | http://emea.support.illumina.com/downloads/local-run-manager-generate-fastq-module.html |  |
| FastQC | http://www.bioinformatics.babraham.ac.uk/projects/fastqc/ |  |
| Bowtie2 v2.3.4.1 | http://bowtie-bio.sourceforge.net/index.shtml |  |
| TopHat v.2.1.1 | <https://ccb.jhu.edu/software/tophat/index.shtml> |  |
| Samtools v1.3 | http://samtools.sourceforge.net |  |
| R | https://www.r-project.org |  |
| EdgeR | <https://bioconductor.org/packages/release/bioc/html/edgeR.html> |  |
| GenomicAlignments | <https://bioconductor.org/packages/release/bioc/html/GenomicAlignments.html> |  |
| GSEA v2.2 | <http://software.broadinstitute.org/gsea/downloads.jsp> |  |
| COMBENEFIT | <https://www.cruk.cam.ac.uk/research-groups/jodrell-group/combenefit> |  |
| Flowing software | <http://flowingsoftware.btk.fi/> |  |
| EMBL | <https://www.embl.de/> |  |
| SPLASHRNA | <http://splashrna.mskcc.org/> |  |
| Proteome Discoverer (PD) 2.4 | ThermoFisher |  |
| SwissProt database | <https://www.uniprot.org/> |  |
| Cytoscape | <https://cytoscape.org/> |  |
| **Instrument** | **Company/Source** |  |
| MyECL™ Imaging System | Invitrogen |  |
| BD FACSCanto II Cell Analyzer | BD Biosciences |  |
| StepOnePlus Real-Time PCR System | ThermoFisher |  |
| Invitrogen Countess II FL Automated Cell Counter | ThermoFisher |  |
| Pannoramic DESK scanner | 3DHISTECH |  |
| FSX100 microscopy | Olympus Life Science |  |
| Fragment Analyzer | Agilent formerly Advanced Analytical |  |
| Axiocam 503 mono + Zeiss axio microscope | Zeiss |  |
| Branson Sonifier 150 | Branson |  |
| Hyrax M55 Rotary Microtome | Leica |  |
| PCR cycler: SimpliAmp thermo cycler | Life technologies |  |
| Orbitrap Fusion Lumos | ThermoFisher |  |
| PCR cycler: SimpliAmp thermo cycler | Life technologies |  |
| Orbitrap Fusion Lumos mass spectrometer | ThermoFisher |  |
| easy-nLC 1200 | ThermoFisher |  |
